# A targeted cell lysis mechanism facilitates toxin release in Clostridioides difficile

**DOI:** 10.1101/2025.09.22.677799

**Authors:** Shannon L. Kordus, Kateryna Nabukhotna, Rubén Cano Rodríguez, Evan Krystofiak, Kevin Childress, Anna J. Smith, Oleg Kovtun, Natalie Loveridge, William Ball, Grace Moore, M. Kay Washington, Nicholas Markham, D. Borden Lacy

## Abstract

*Clostridioides difficile* infection depends on the production of two large toxins, TcdA and TcdB, encoded within a pathogenicity locus alongside the phage-like holin TcdE. The mechanism of toxin secretion remains actively debated, with current models proposing either TcdE-dependent non-lytic secretion or TcdE-independent lytic release. Here, we provide evidence for a unifying model where TcdE drives lysis in a phenotypically distinct subpopulation of cells. We show that TcdE, TcdA, and TcdB expression is restricted to a small fraction of cells exhibiting markers of active lysis and establish this subpopulation as the driver of severe pathogenic outcomes in a mouse model of CDI. Overexpression of TcdR, the sigma factor regulating the pathogenicity locus, triggers TcdE-dependent lysis, even in strains previously reported to employ TcdE-independent secretion mechanisms. Correlative light and electron microscopy combined with cryo-ET reveal a distinctive ultrastructure in lytic cells. Membrane vesicles accumulate between a disrupted inner membrane and intact peptidoglycan, alongside an electron-dense material containing TcdA. These observations reveal a population-level strategy in which a minority of bacteria sacrifice themselves through TcdE-mediated lysis to release toxins as a ‘bet-hedging strategy’.

## Introduction

*C. difficile* is a Gram-positive, spore-forming, anaerobic bacterium that infects the colon to cause a range of disease severity from mild cramps and diarrhea to more severe, life-threatening sequelae. The greatest population at risk for *C. difficile* infection (CDI) are people over 65 years of age who have been treated with broad spectrum antibiotics, are immunosuppressed, or have other chronic comorbidities; however, infections in young adults are now common^1,2^. The primary virulence factors associated with CDI are two toxins, TcdA and TcdB. The toxins are large proteins that cause loss of cellular junctions, inflammation, and cell death^3,4^. While much is known regarding toxin effects on the host, the mechanism(s) by which the toxins are released from the bacterium remains unclear.

Bacterial protein secretion typically relies on N-terminal secretion signals to direct cargo to dedicated export systems^5^. However, TcdA and TcdB lack these canonical targeting sequences, suggesting an alternative release mechanism. Two competing mechanisms on TcdA and TcdB release have been proposed: **(1)** lytic release during the bacterial lifecycle^6,7^ and **(2)** non-lytic release mediated by TcdE, a phage-like holin that permeabilizes the inner membrane^8–12^.

Bacteriophages promote population-level bacterial lysis through the coordinated action of holins and endolysins^13–15^. Holins form pores in the inner membrane^13,16,17^, allowing endolysins to access the periplasm and degrade peptidoglycan. In Gram-positive bacteria, holins and endolysins are sufficient for lysis^18^. While in Gram-negative bacteria, spanins are needed for fusing the inner and outer membranes or permeabilizing the outer membrane^14,19^. Phage lysis cassettes (PLCs) consisting of a holin, endolysin, and, in Gram negative bacteria, spanin, have been identified in various bacteria where they are encoded alongside virulence factor genes and have been designated Type 10 secretion systems in some studies^9,20–27^.

The *C. difficile* pathogenicity locus encodes the toxins TcdA and TcdB, transcriptional regulators TcdR (positive) and TcdC (negative), and two bacteriophage-homologous genes, *tcdE* and *tcdL*. TcdE shares high sequence homology to λ phage holin S, while TcdL is a truncated, catalytically inactive homolog of λ phage endolysin R. Conservation of both *tcdE* and *tcdL* across all toxin-producing *C. difficile* strains suggests a functional role in toxin release^9^. However, the contribution of TcdE to toxin release remains unclear. In some strains, *tcdE* deletion did not significantly reduce toxin release^6^, whereas in others, deletion substantially decreased toxin released *in vitro* and disease severity *in vivo*^10,28,29^. These conflicting results have led to competing mechanistic proposals, with most studies suggesting TcdE functions through lysis-independent pathways.

The conflicting mechanistic models underscore a fundamental gap in our understanding of toxin release. To resolve these controversies, we systematically examined toxin release dynamics in multiple strains using high-sensitivity detection methods and direct visualization of cellular changes accompanying toxin release. We find that TcdE expression is restricted to a small subpopulation of cells that undergo lysis, and that this population-level strategy is sufficient to drive pathogenic outcomes in vivo. Our results reconcile conflicting studies and establish TcdE-mediated lysis as the primary route of toxin release in *C. difficile* infection.

## Results

### Toxin release varies across *C. difficile* strains

To investigate the mechanism of toxin release in *C. difficile*, we sought to address several outstanding questions. Does the timing and production level of TcdA and TcdB differ across strains? Is toxin release coordinated with synthesis, or do bacteria accumulate intracellular toxin that is released asynchronously from production? Does toxin release correlate with bacterial lysis? And given prior reports of strain-specific mechanistic differences, which observations are generalizable across strains versus strain-dependent?

Using newly developed nanobody-based ELISAs capable of detecting femtomolar concentrations of TcdA and TcdB^30^, our first goal was to quantify intracellular and extracellular TcdA and TcdB levels throughout *in vitro* bacterial growth. We selected four strains that represent a spectrum of *C. difficile* virulence and toxin production profiles. The laboratory strain 630Δerm (ribotype (RT) 012) causes mild infection and produces minimal toxin, with prior evidence indicating TcdE-independent toxin release^6,31,32^. In contrast, VPI10463 (RT087) produces high levels of TcdB and causes severe disease outcomes in animal models^31^. The epidemic-associated RT027 strains R20291 and M7404 display intermediate phenotypes and toxin levels relative to these extremes^31^. This strain panel thus allows us to examine toxin dynamics across a clinically relevant range of virulence levels and to evaluate whether release mechanisms are conserved or strain-specific.

Bacteria were grown in nutrient-limiting medium and sampled at 4, 8, and 18 hours to capture log-, stationary-, and death phases, respectively. Nanobody-based ELISAs were used to measure toxin concentrations in the bacterial supernatant and pellet or the total amount in the bacterial culture. Colony forming units (CFUs) were quantified at each timepoint to assess growth phase and ensure similar amounts of bacteria were present when quantifying toxin in supernatants. All four strains exhibited similar growth kinetics, with comparable CFU counts at each timepoint (Figure 1, lines).

**Figure 1.**
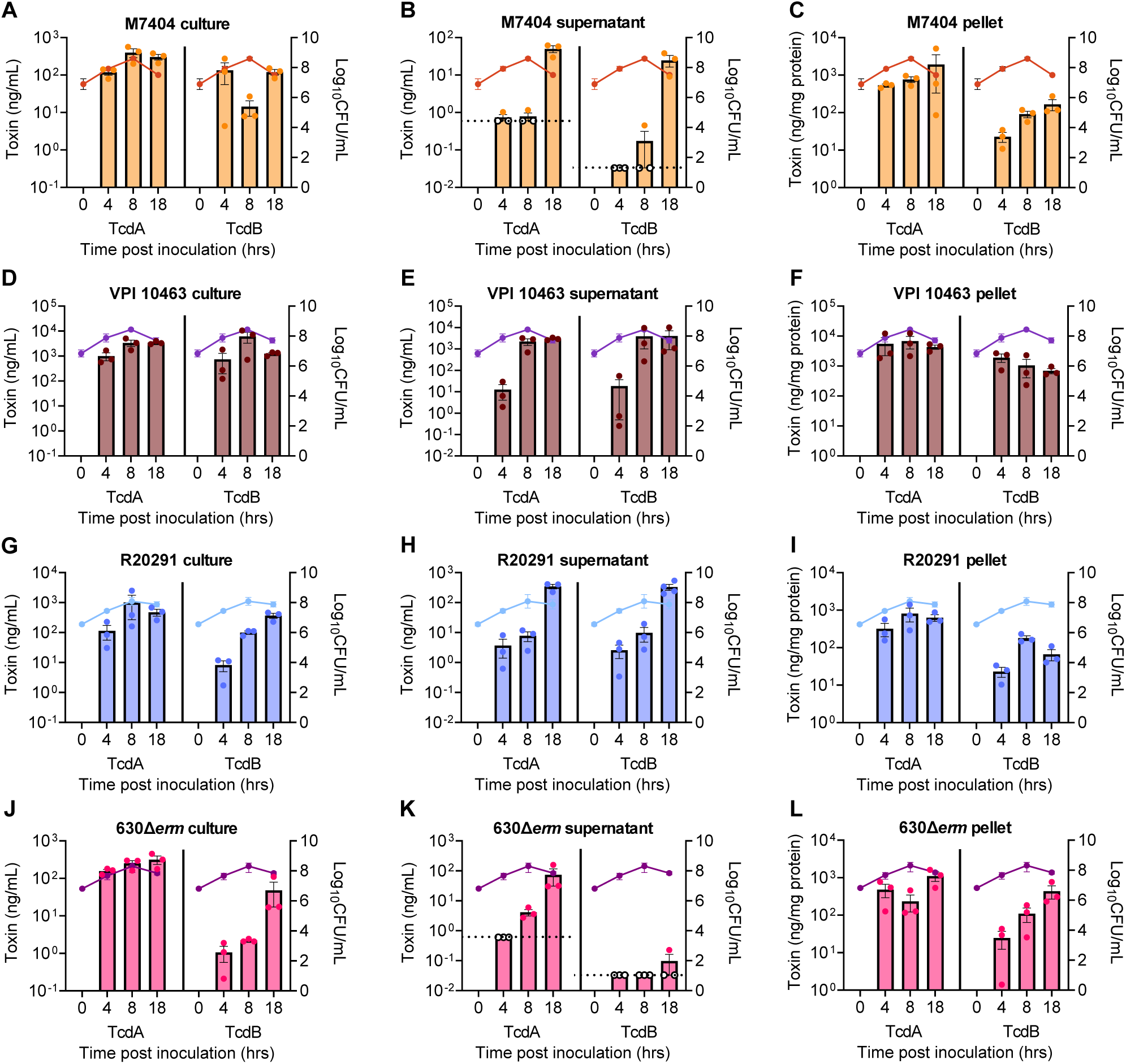
A comparison of toxin release between different *C. difficile* strains reveals two patterns of toxin release. Bacterial cultures were grown in TY medium and at various time points TcdA and TcdB was measured by ELISA. Pink line represents the colony forming units (CFUs) at each time point. **A)** Culture, (**B)** Supernatant, and **(C)** Pellet TcdA and TcdB measured in M7404 at 4 hrs (mid log phase), 8 hrs (early stationary phase), and 18 hrs (death phase) time post inoculation. **D)** Culture, **(E)** Supernatant, and **(F)** Pellet TcdA and TcdB measured in VPI 10463 at 4, 8, and 18hrs time post inoculation. **G)** Culture, **(H)** Supernatant, and **(I)** Pellet TcdA and TcdB measured in R20291 at 4, 8, and 18hrs time post inoculation. **J)** Culture, **(K)** Supernatant, and **(L)** Pellet TcdA and TcdB measured in 630Δ*erm* at 4, 8, and 18hrs time post inoculation. Empty circles represent ELISA values that fell below the limit of detection, dotted line. Each data point is the average of two technical replicates. Error bars represent SEM.

Despite having similar growth kinetics, two patterns of toxin release were observed: VPI10463 and R20291 released TcdA and TcdB continuously throughout all growth phases, with detectable extracellular toxin present at 4 hours and increasing through stationary phase (Figure 1E,H). In contrast, 630Δ*erm* and M7404 displayed predominantly death phase toxin release, with little to no detectable extracellular toxin at 4 and 8 hours despite substantial intracellular accumulation at these same timepoints (Figure 1A-C, J-L).

Examination of intracellular toxin levels revealed that all strains accumulated detectable TcdA and TcdB in bacterial pellets by the 4-hour timepoint (Figure 1C,F,I,L). TcdA levels remained relatively stable throughout the growth curve in all strains, suggesting constitutive production followed by either release or retention. In contrast, TcdB accumulated over time in M7404 and 630Δ*erm*, reaching higher intracellular concentrations by 18 hours compared to earlier timepoints (Figure 1A,C,J,L).

A striking observation was that intracellular TcdA and TcdB concentrations in early-phase cultures (4–8 hours) of 630Δerm were comparable to those in the epidemic strains M7404 and R20291 (Figure 1A,C,G,I,J,L). Yet 630Δ*erm* released minimal toxin at these early-time points despite harboring comparable intracellular concentrations. This was also observed when comparing the intracellular concentrations of TcdA and TcdB between VPI10463 and R20291. VPI10463 had nearly 10-fold more intracellular TcdA and TcdB concentrations than R20291 yet, in log phase (4 hrs), both strains released similar amounts of toxins (Figure 1D-I). The maintenance of relatively constant intracellular toxin levels throughout the growth curve, despite ongoing release in early-release strains, further demonstrates that toxin synthesis and release are uncoupled temporal processes. Strain-specific differences in virulence thus correlate more closely with the *timing* and *efficiency* of toxin secretion than with the absolute amount of toxin produced.

### TcdE influences toxin release *in vitro* in a strain-dependent manner

To determine the role of TcdE in toxin release, we next created isogenic *tcdE* deletions in R20291, an early-release strain, or in 630Δ*erm,* a death phase-release strain. In R20291Δ*tcdE,* TcdA and TcdB were largely undetectable in the supernatant during the log and stationary phases, (Figure 2A-B, dotted line and empty circles, Extended Data Figure 1B,E) despite intracellular and total toxin concentrations comparable to wild-type R20291 (Figure 2C-D; Extended Data Figure 1A-F). At death-phase, R20291Δ*tcdE* released significantly less toxin than R20291 (Figure 2A-B; Extended Data Figure 1B,E). In contrast, the 630Δ*erm*Δ*tcdE* strain showed only a modest reduction in TcdA release during stationary and death phases, with no significant difference in TcdB release despite harboring similar intracellular and total toxin levels as the parent strain (Figure 2A-D; Extended Data Figure 1G-L). These results align with prior reports indicating TcdE-dependent toxin release in R20291 but TcdE-independent mechanisms in 630Δ*erm*^6,11^.

**Figure 2.**
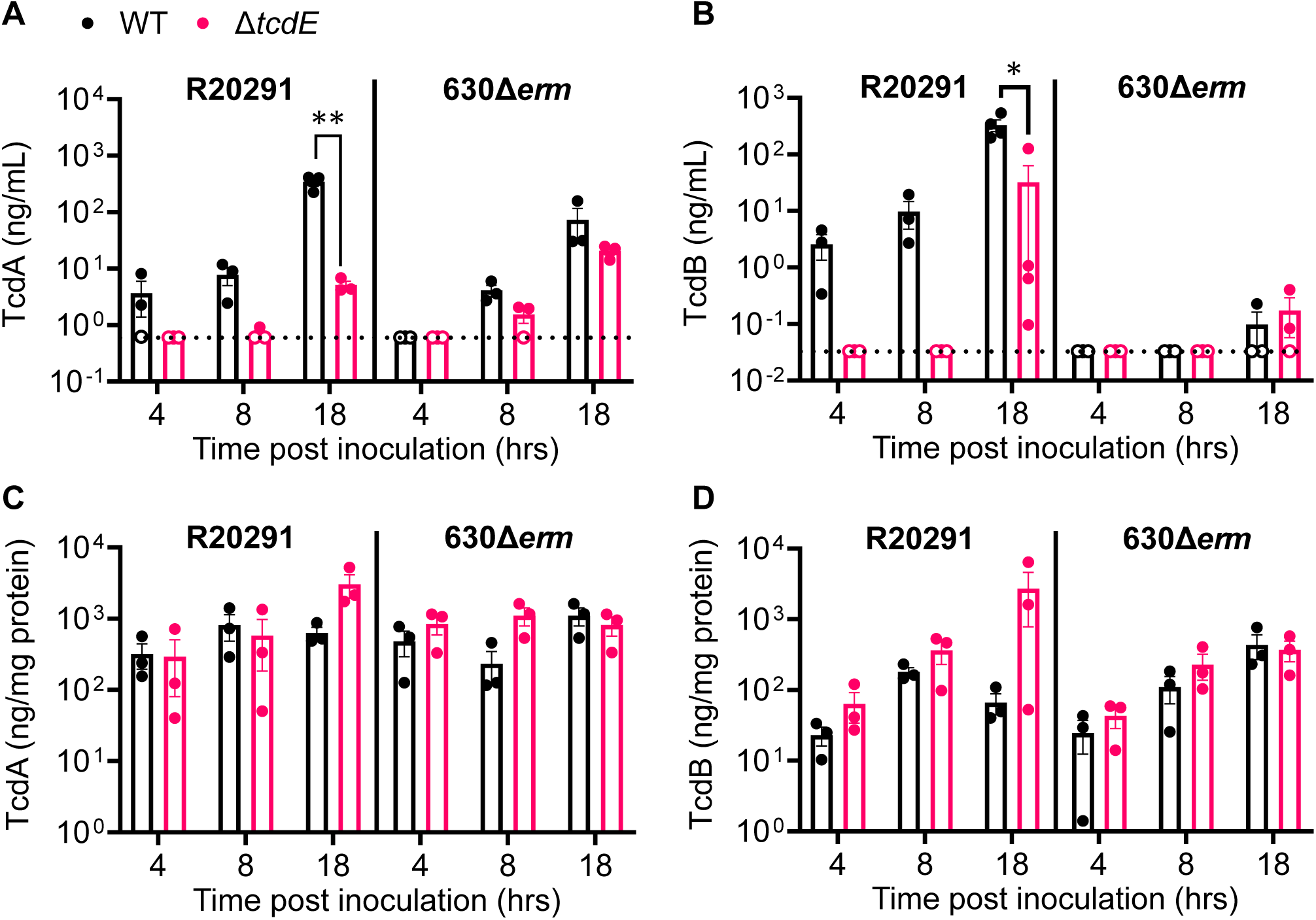
Assessing the role of *tcdE* on toxin release in *C. difficile*. Bacterial cultures were grown in TY medium and at various time points, the TcdA and TcdB in culture supernatants were measured by ELISA. Supernatant **(A)** TcdA or **(B)** TcdB were measured in R20291 (black, left), R20291Δ*tcdE* (pink, left), 630Δ*erm* (black, right), and 630Δ*erm*Δ*tcdE* (pink, right) at (mid-log phase), 8 hrs (early stationary phase), and 18 hrs (death phase) post inoculation. **(C)** TcdA or **(D)** TcdB concentrations within the bacterial pellet were measured in R20291 (black, left), R20291Δ*tcdE* (pink, left), 630Δ*erm* (black, right), and 630Δ*erm*Δ*tcdE* (pink, right) at the same time points. Data in C and D are normalized to total protein. Empty circles represent ELISA values that fell below the limit of detection, dotted line. Each data point is the average of two technical replicates. Data shown as mean and error bars as SEM. A Sidak unpaired t-test was used to calculate statistical significance (** *P*<0.01, * *P*<0.05).

### TcdE influences virulence through toxin release *in vivo* in a strain-dependent manner

To determine if the strain-specific differences in TcdE-dependent timing and/or efficiency of toxin release observed *in vitro* also correlate with virulence *in vivo*, we used a mouse cefoperazone CDI model^31^ to compare disease severity in mice infected with R20291 or R20291Δ*tcdE*. Deletion of *tcdE* in R20291 resulted in attenuated weight-loss throughout infection (Figure 3A, Extended Data Figure 2A-D) and significantly reduced cecal edema, colonic inflammation, and colonic epithelial injury at day 4 post infection (Extended Data Figure 2E-H).

**Figure 3.**
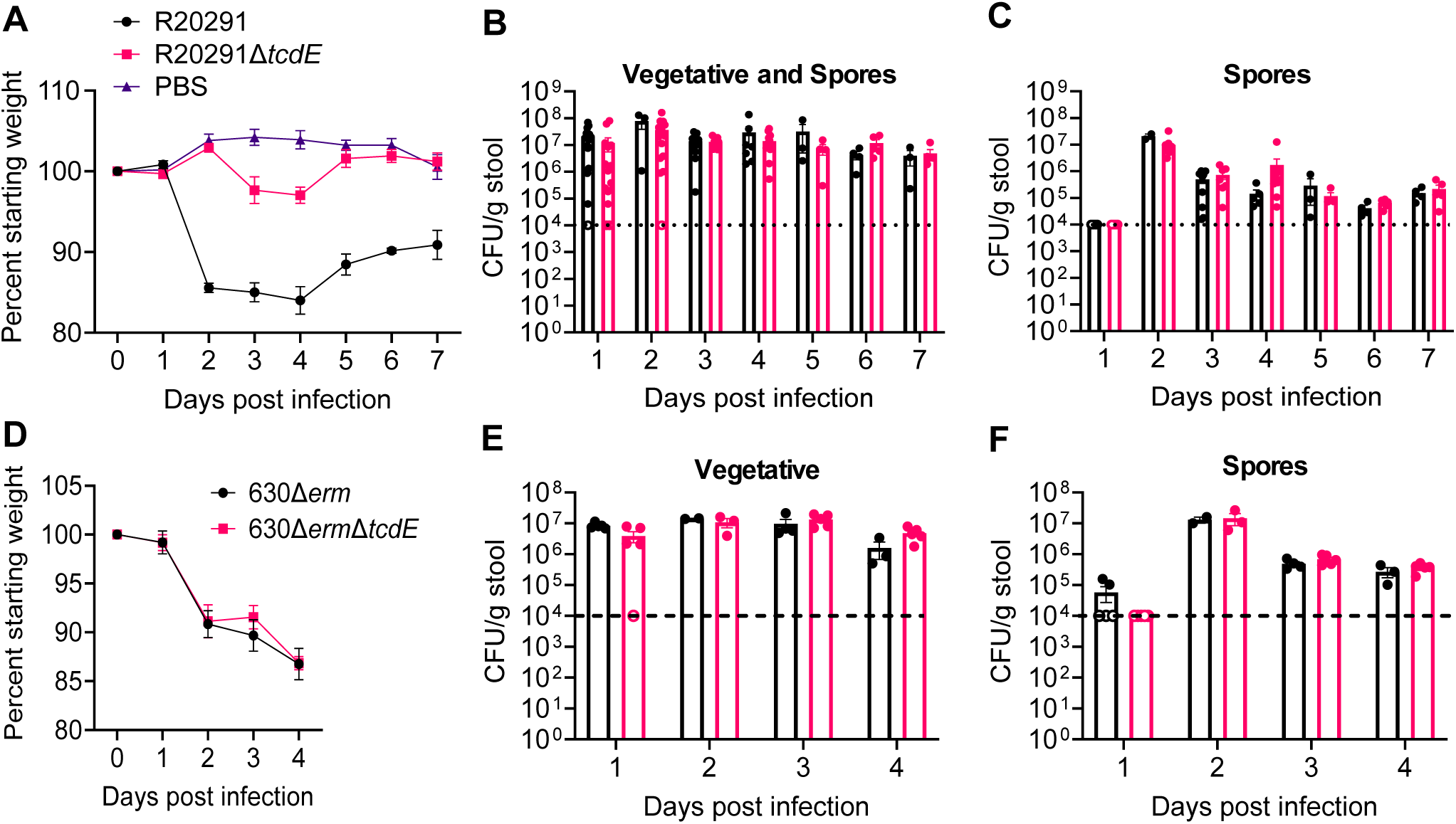
TcdE-dependent toxin release causes less severe disease *in vivo* despite similar colonization. Mice were infected with 100 spores of R20291, R20291Δ*tcdE*, or a PBS vehicle control, and **(A)** weight loss was tracked for 7 days. Stool was collected every day for 7 days and plated for **(B)** vegetative bacteria and spores or **(C)** spores alone. **D)** Mice were infected with 100 spores of 630Δ*erm* or 630Δ*erm*Δ*tcdE,* and weight loss was tracked for 4 days. Stool was collected every day for 4 days and plated for **(E)** vegetative bacteria or **(F)** spores. Data shown as mean and error bars as SEM. Dotted line represents limit of detection, samples that are empty circles represent values that fall below limit of detection.

Bacterial burdens were comparable between R20291 and R20291Δ*tcdE* throughout the infection with no significant differences in the levels of total bacteria (vegetative bacteria and spores) or spores alone (Figure 3B-C), indicating that *tcdE* deletion did not impair colonization. To determine whether disease attenuation was correlated with toxin levels, we measured TcdA and TcdB in stool samples (which were freeze-thawed, reflecting both intracellular and extracellular toxin). On day 1 post infection, neither strain produced detectable toxin in most mice, though some R20291 infected mice had measurable TcdA (Extended Data Figure 3A-B; dotted line and empty circles). By day 2, R20291Δ*tcdE*-infected mice had significantly higher TcdA levels than R20291-infected mice (Extended Data Figure 3A). TcdB concentrations were similar between strains throughout the infection (Extended Data Figure 3B).

In stark contrast, *tcdE* deletion in 630Δ*erm* had minimal impact on disease outcomes. Mice infected with 630Δ*erm* and 630Δ*erm*Δ*tcdE* exhibited similar weight-loss, survival (Figure 3D, Extended Data Figure 4A-C), and comparable histopathological scores for edema, inflammation, and epithelial injury in the cecum and colon on day 4 post infection (Extended Data Figure 4D-G). Both strains achieved similar colonization levels (Figure 3E-F) and produced comparable concentrations of TcdA and TcdB during infection (Extended Data Figure 3-B). These findings demonstrate that the strain-specific role of TcdE in toxin release, observed *in vitro*, translates to reduced virulence *in vivo*.

### TcdE contributes to *C. difficile* cell lysis

To visualize TcdE-dependent toxin release, we engineered R20291 to express a TcdE-mScarlet fusion within the native pathogenicity locus. These cells were counterstained with the lipophilic, amine-reactive membrane stain CellBrite 488. Imaging of 779 bacteria revealed that ∼16% expressed TcdE as a small punctum localized to the cell membrane (Figure 4A). Notably, the ∼0.4% of cells with multiple TcdE puncta also displayed high CellBrite 488 accumulation, suggesting membrane permeabilization or disruption (Figure 4A, inset).

**Figure 4.**
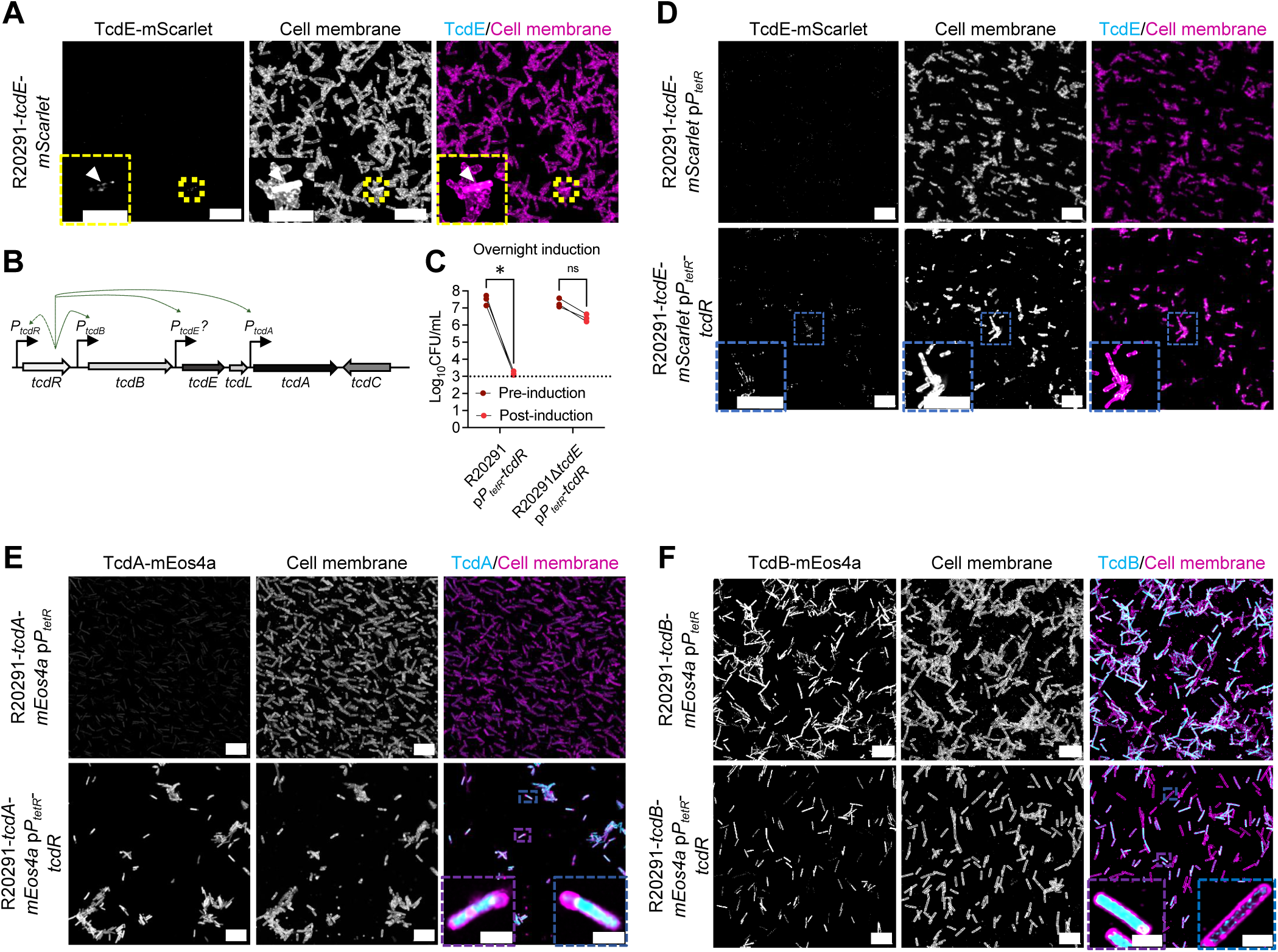
TcdE drives lysis in a small population of bacteria expressing toxins. **A)** Maximum intensity projection lightening confocal image of fixed R20291 showing TcdE (cyan) and cell membrane (magenta) (scale bar 10 µm). Inset of yellow box in top panels. Arrow points to cell expressing TcdE (scale bar 5 µm). **B)** Pathogenicity locus highlighting regulatory elements of the positive regulator TcdR. **C)** Viability time course comparing R20291 or R20291Δ*tcdE* before (dark red) or after (light red) *P_tetR_* induction. Induction of *P_tetR_* began at 8 hrs post inoculation for 16 hrs. A 2-way-ANOVA followed by an Uncorrected Fisher’s LSD post hoc test was used to test for significance. Significance is reported as * *P*<0.05. D) Maximum intensity projection lightening confocal image of fixed R20291 showing TcdE (cyan) and cell membrane (magenta) empty vector control or with induction of *tcdR* by *P_tetR_* (scale bar 10 µm). Induction by *P_tetR_* occurred after 18 hrs of growth for 4 hrs. Inset box points to cells expressing high amounts of TcdE colocalizing with membrane (scale bar 5 µm). **E)** Maximum intensity projection lightening confocal image of fixed R20291 showing TcdA (cyan) and cell membrane (magenta) empty vector control or with induction of *tcdR* by *P_tetR_* (scale bar 10 µm). Induction by *P_tetR_* occurred after 18 hrs of growth for 4 hrs. Blue inset box points to cell expressing high amounts of TcdA but no co-localization with membrane (scale bar 2 µm). **F)** Maximum intensity projection lightening confocal image of fixed R20291 showing TcdB (cyan) and cell membrane (magenta) empty vector control or with induction of *tcdR* by *P_tetR_* (scale bar 10 µm). Induction by *P_tetR_* occurred after 18 hrs of growth for 4 hrs. Blue inset box points to cell expressing high amounts of TcdB but no co-localization with membrane (purple inset) or low amounts of TcdB (blue inset) (scale bar 2 µm). Data shown as mean and error bars as SEM.

To overcome the scarcity of TcdE-expressing cells, we overexpressed *tcdR* using an anhydrotetracycline (ATc)-inducible promoter (p*P_tetR-_tcdR*). Since TcdR regulates *tcdA* and *tcdB* expression and is predicted to drive expression of *tcdE* (Figure 4B)^29,33^, we hypothesized that this would increase TcdE levels. Indeed, *tcdR* induction substantially increased TcdE-mScarlet expression compared to the empty vector controls (Figure 4D). Consistent with native expression, high membrane stain colocalized with multiple TcdE puncta, supporting the hypothesis that elevated TcdE expression compromises membrane integrity. Further, when incubated overnight we noted a signficant reduction in CFUs following TcdR induction (Figure 4C).

To test whether TcdR-dependent lysis is mediated by TcdE, we introduced p*P_tetR_* or p*P_tetR-_tcdR* into R20291, 630Δ*erm,* and their respective *tcdE* deletion strains. Uninduced strains showed no growth differences when comparing empty vector and *tcdR* constructs (Extended Data Figure 6A-B; 7A-B). Upon *tcdR* induction, R20291 displayed a ∼2-fold reduction in doubling time (1.8±0.8 hrs vs. 3.8±0.5 hrs) and significantly reduced vegetative viability at 6 and 8 hours that did not recover to control levels (Extended Data Figure 6E–F). Notably, spore numbers remained unchanged, indicating lysis rather than sporulation (Extended Data Figure 6F, H). 630Δ*erm* showed even more dramatic effects: *tcdR* induction resulted in complete loss of viability by 24 hours and significant vegetative reduction at 4, 6, and 8 hours (Extended Data Figure 7E–F). Critically, *tcdE* deletion in R20291 and 630Δ*erm* abolished the *tcdR*-induced lysis phenotype in both strains (Extended Data Figures 6G-H and 7G-H). R20291Δ*tcdE* showed no significant reduction in doubling time upon *tcdR* induction (1.7±0.8 hrs vs. 1.4±0.6 hrs; Extended Data Figure 6G) and recovered viability to control levels (Extended Data Figure 6H). Similarly, 630Δ*erm*Δ*tcdE* maintained viability at 24 hours, unlike the parent strain (Extended Data Figure 7H). In stationary phase cultures, R20291 p*P_tetR-_tcdR* showed a ∼10,000-fold loss in viability after 16 hours, while R20291Δ*tcdE* p*P_tetR-_tcdR* showed no change (Figure 4C). These data demonstrate that TcdE mediates TcdR-induced cell death even in strains that did not show TcdE-dependent toxin release.

To determine whether toxin accumulates in lysing cells, we imaged TcdA-mEos4a and TcdB-mEos4a fusion proteins. Empty vector controls showed low, inconsistent TcdA signal, while TcdB signal was variable across the population (Figure 4E–F). Upon *tcdR* induction, both toxins increased in R20291 (Figure 4E–F), but high CellBrite 640 membrane stain did not consistently overlap with high toxin levels. Importantly, cells expressing high TcdA or TcdB showed concentrated cytoplasmic signal (Figure 4E–F, inset). This suggests that TcdA and TcdB are not uniformly expressed in a large population of cells, but likely in a sub-population that also express high amounts of TcdE.

### Cells expressing *tcdR* and *tcdE* reveal intracellular vesicle formation and cellular lysis

To further understand how TcdE contributes to cellular lysis, we used complementary high-resolution electron microscopy approaches to visualize cellular ultrastructure. First, using freeze-substituted transmission electron microscopy (FS-TEM), we compared R20291 and R20291Δ*tcdE* overexpressing TcdR to their respective empty vector controls. In both empty vector controls and R20291Δ*tcdE* overexpressing TcdR, the bacterial envelope appeared intact with well-resolved inner membrane, thick peptidoglycan, and a paracrystalline surface layer (S-layer) (Extended Data Figure 8A-B). In contrast, R20291 overexpressing TcdR displayed striking ultrastructural changes: vesicles protruded from the inner membrane into the periplasmic space, the inner membrane appeared discontinuous, and an electron dense material was observed at sites of membrane disruption (Extended Data Figure 8A-C).

To correlate toxin localization with these observations, we used cryo-electron tomography (cryo-ET) to visualize R20291-*tcdA-mEos4a* overexpressing *tcdR*. Cells were imaged by cryo-confocal and thinned using cryo-focused ion beam-scanning electron microscopy (cryo-FIB-SEM) to create lamellas. By correlating the fluorescent images from the cryo-confocal with the EM-search maps, we collected tomograms on bacteria with and without TcdA-mEos4a expression. Bacteria expressing low levels of TcdA displayed normal ultrastructure with an intact inner membrane, peptidoglycan, and S-layer (Figure 5A-E, Supplementary Movie 1). In contrast, bacteria with high TcdA expression exhibited severe membrane disruption: the inner membrane detached from the peptidoglycan with visible loss of continuity and vesicle formation, while the S-layer became discontinuous and the peptidoglycan remained largely intact (Figure 5F-J, Supplementary Movie 2). Notably, the electron dense regions at the sites of membrane damage remained apparent and correlated with the sites of TcdA localization.

**Figure 5.**
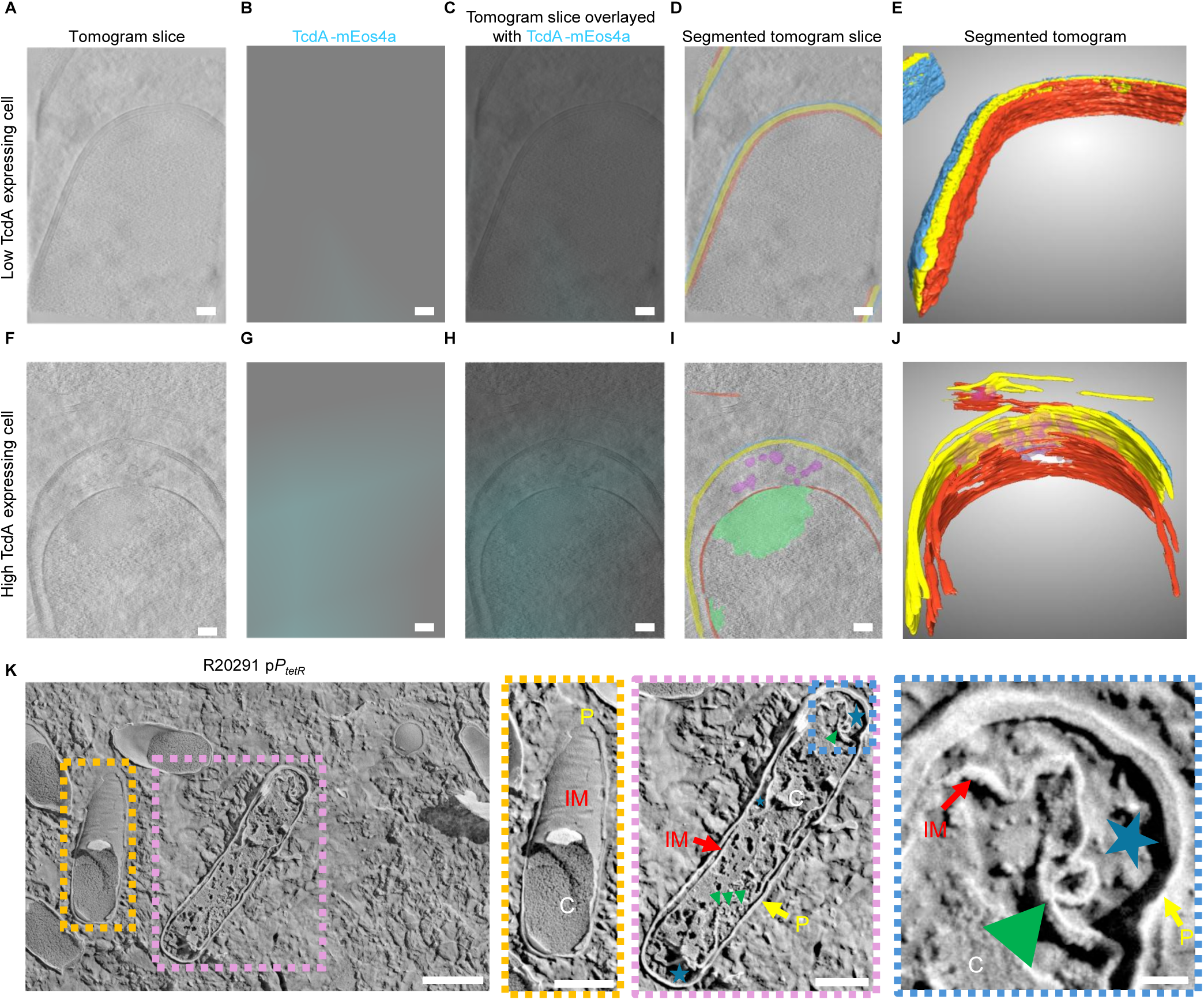
Cryogenic ultrastructure of bacteria expressing high and low amounts of TcdA. **A)** A single tomogram slice from R20291-*tcdA-mEos4* p*P_tetR_-tcdR* representing low TcdA expressing bacteria. **B)** Cryo-confocal image representing TcdA-mEos4a expression in **(A)**. **C)** Overlay of **(A)** and **(B)**. **D)** Segmented tomogram overviews represent inner membrane, red; peptidoglycan, yellow. **E)** Fully segmented 3D tomogram from **(A)**. **F)** A single tomogram slice from R20291-*tcdA-mEos4* p*P_tetR_-tcdR* representing high TcdA expressing bacteria. **G)** Cryo-confocal image representing TcdA-mEos4a expression in **(F)**. **H)** Overlay of **(F)** and **(G)**. **I)** Segmented tomogram overviews represent inner membrane, red; peptidoglycan, yellow; S-layer, blue; electron dense region, green; lipid bilayer vesicles, purple. **J)** Fully segmented 3D tomogram from **(H)**. Scale bar 100 nm. **K)** Freeze fractured R20291 p*P_tetR_* highlighting a “healthy” bacterium (orange inset) and pre-lytic bacterium (light purple and blue inset). C, cytoplasm (white); IM, inner membrane (red); P, peptidoglycan (yellow). Green triangles indicate vesicles and blue stars indicate inner membrane and peptidoglycan detachment. Scale bar 1 μm, blue inset 200 nm.

To exclude the possibility that these ultrastructural changes resulted from TcdR overexpression artifacts, we examined R20291 with the empty vector using freeze-fracture cryo-SEM. Most bacteria displayed intact inner membranes with continuous peptidoglycan layers and cytoplasm (Figure 5K, orange inset). However, we also observed a subpopulation of R20291 cells with detached inner membranes, periplasmic vesicles, and discontinuous cytoplasm (Figure 5K, light purple and blue inset), indicating that this phenotype occurs naturally and is not an artifact of overexpression.

To determine if the vesicles contained toxins, we purified vesicles from supernatants of R20291, R20291Δ*tcdE*, and R20291 *tcdAB::CT* (ClosTron functional disruptions in *tcdA* and *tcdB*) using iodixanol gradient centrifugation. Negative stain EM revealed extracellular vesicles in fractions 6 and 9 of R20291, whereas fraction 9 of R20291Δ*tcdE* contained phage- or difficin-like complexes instead of vesicles (Extended Data Figure 9A-C). In contrast, R20291 *tcdAB::CT* retained vesicles in fraction 9 (Extended Data Figure 13C). Toxin detection by ELISA showed that TcdA and TcdB were enriched in fraction 9 of R20291 but absent from R20291Δ*tcdE* or R20291 *tcdAB::CT*. To test whether the toxins were vesicle associated, we applied fraction 9 from each strain to Vero-GFP cells. Only R20291 fraction 9 induced cell rounding, which was neutralized by anti-TcdA/B antibodies (Extended Data Figure 13D-E). This result indicates that while TcdA and TcdB co-fractionate with vesicles, they are not encapsulated within the vesicle. Collectively, these data suggest that the intracellular vesicles observed by EM are a direct consequence of TcdE-mediated membrane disruption but are not the primary vehicle for toxin release.

## Discussion

In this study, we show that the Gram-positive pathogen *C. difficile* utilizes a phage holin-like protein to release TcdA and TcdB through specialized cellular lysis in a small subset of the bacterial population to cause disease. A recent report from Dupuy and colleagues was similarly focused on the impact of TcdE on TcdA and TcdB release^10^. Our study provides independent validation of several key points: (1) Deletion of *tcdE* does not impact the intracellular levels of TcdA and TcdB. (2) Deletion of *tcdE* in a 027 strain (e.g., UK1 or R20291) reduces extracellular TcdA and TcdB release *in vitro* and causes loss of virulence *in vivo* without impacting bacterial fitness. (3) Using two different mouse models of *C. difficile* infection (gnotobiotic or cefoperazone-treated), our studies indicate that TcdE contributes to toxin release *in vivo* and that alternative mechanisms such as cell death or sporulation, do not appear to be significant drivers of virulence.

The major conceptual advance in this work is the interpretation of how TcdE-dependent toxin release occurs. Previous frameworks treated bacterial cultures as a homogenous population, using population-level assays (e.g., CFU/optical density, lactate dehydrogenase assays and flow cytometry with propidium iodide staining) to conclude that TcdE-mediated release occurs independently of lysis^10–12^. However, if TcdE-dependent lysis occurs only in a small subset of cells, population-level measurements would mask these single-cell phenotypes.

To visualize TcdE function at the single-cell level, we created a *C. difficile* strain expressing a TcdE-mScarlet fusion. Quantification revealed that ∼16% of cells expressed low levels of TcdE and ∼0.6% expressed high amounts. Cells with high TcdE expression coincided with high intracellular accumulation of the membrane dye CellBrite, suggesting compromised membrane integrity.

Because only ∼0.6% of cells were lysing, population-level assays would mask this phenotype. Therefore, we overexpressed the sigma factor TcdR, which positively regulates *tcdA*, *tcdB*, and its own expression^33,34^. TcdR overexpression increased TcdE-mScarlet fluorescence, indicating that TcdR also positively regulates *tcdE*. Importantly, TcdR overexpression decreased vegetative viability in R20291 and 630Δ*erm* without increasing sporulation. Deletion of *tcdE* in both strains restored viability, confirming a role for TcdE in lysis.

The artificial overexpression of TcdE, TcdA, and TcdB through TcdR induction promotes TcdE-dependent cell death even in 630Δ*erm*, a strain previously thought to exhibit TcdE-independent toxin secretion^6^. Based on these findings, we propose a new model. The total amount of TcdA and TcdB made by the bacterial population is not the sole driver of disease; instead, disease is driven by TcdR expression of the *PaLoc* within a cellular subset. The lysis of these cells then releases the virulence factors necessary to cause disease. This differentiated cell program sacrifices a small population to benefit the remaining community, a phenomenon known as bacterial bet-hedging^35^. We propose that toxin action on the host generates “communal goods” or access to new ecological niches that benefit the broader population.

This model is strongly supported by several key reports from the literature. First, heterogeneous toxin production within *C. difficile* populations has been demonstrated and is controlled by a stochastic, phase-variable, riboswitch^36–39^. Second, holin-dependent lysis and virulence factor release has been elegantly demonstrated in the Gram-negative organisms *Yersinia entomophaga* and *Serratia marcescens*^21,22^. In these systems, the virulence factor expression in a subset of cells is coordinated with a ‘phage lysis cassette’ containing holin, endolysin, and spanin. The holin destabilizes the inner membrane, allowing endolysin access to degrade the peptidoglycan layer. Spanins then fuse the inner and outer membranes to promote release of intracellular contents and vesicles. Notably, the holin-formed pores do not directly translocate virulence factors; rather, they facilitate endolysin access to the cell wall and spanin(s) access to the membranes.

A persistent challenge in understanding *C. difficile* toxin release has been reconciling observations between high- and low-toxin producing strains. We observed that two high-toxin-producing strains (VPI10463 and R20291) release toxins in a TcdE-dependent manner, while 630Δ*erm* appeared TcdE-independent. Surprisingly, despite its reputation as a low toxin producer, 630Δ*erm* accumulates TcdA and TcdB intracellularly at levels comparable to R20291. We propose that low extracellular toxin from 630Δ*erm* results not from low toxin production, but from low TcdE expression. Supporting this hypothesis, Mehner-Breitfeld e*t*. *al*. identified differences in the putative ribosome binding sites upstream of *tcdE* between 630Δ*erm* and high-toxin producing strains^9^. Additionally, strain-dependent phase-variable riboswitches have been reported: in 630Δ*erm* the riboswitch is locked in the OFF-orientation, while in R20291 it can adopt both ON and OFF states^36,39^. This mechanism likely explains low toxin release from 630Δ*erm* and the lack of phenotype when mice were challenged with 630Δ*erm*Δ*tcdE*.

Canonical bacteriophage lysis in Gram-positive bacteria requires a functional endolysin to degrade peptidoglycan. Directly downstream of *tcdE* lies a predicted partial endolysin, *tcdL,* conserved across all toxin-producing *C. difficile* strains but lacking catalytic activity. This deficit has prompted investigations of peptidoglycan remodeling enzymes outside of the *PaLoc*. Two studies identified such candidates: a lytic-transglycosylase, Cwp19, which induces autolysis under specific conditions^40^ and an endolysin capable of releasing TcdA independent of population level lysis^8^. The role of these proteins at the single cell level remains unknown. Our study focused on TcdE dependence without investigating endolysin contributions, although our EM data showed that the peptidoglycan was largely intact. Further work is needed to understand peptidoglycan remodeling during *C. difficile* lysis.

Our study is the first to demonstrate that *C. difficile* toxin release occurs through holin-dependent lysis of a small bacterial subpopulation, a mechanism that mirrors Gram negative systems. Holin and/or endolysin mediated mechanisms are believed to allow certain toxins, such as typhoid toxin and those from *C. perfringens* and *P. sordellii*, to be released without cellular lysis^24,26,27^. However, further studies are warranted to investigate this process at a single cell level. The data presented here support the growing body of literature demonstrating that bacteria can co-opt a phage lysis cassette to coordinate the expression and release of virulence factors in a small subset of cells to drive disease progression. The environmental factors that stimulate this differential programming both within and across strains warrants further investigation.

## Methods

### Bacterial growth conditions, medium, and strains

*Clostridioides difficile* strains were grown in a strict anaerobic environment within a COY anaerobic chamber (5% H_2_, 5% N_2_, and 90% CO_2_) statically at 37°C. Strains were grown in BHIS (brain heart infusion-supplemented with 5 g/L yeast extract) medium, TY (30 g/L tryptone and 20 g/L yeast extract) medium, or CDDM^41^ supplemented with 0.1% L-cysteine. For toxin release assays, L-cysteine was omitted from the medium. *C*. *difficile* growth medium was supplemented with the following when needed: 0.1% taurocholate [TA], cefoxitin (8 μg/mL), thiamphenicol [Thi] (10 μg/mL or 15 μg/mL), kanamycin [Kan] (50 μg/mL), D-cycloserine (250 μg/mL), lincomycin (20 μg/mL), anhydrous tetracycline [ATc] (20 ng/mL), uracil (5 µg/mL), 5-Fluoroorotic Acid [5-FOA] (5 µg/mL) or 1% (w/v) D-xylose. *Escherichia coli*, *Bacillus megaterium, Bacillus subtilis* strains were maintained on Lysogeny Broth (LB) at 37°C supplemented with either erythromycin [Erm] (200 μg/mL), chloramphenicol [Cam] (34 μg/mL for *E*. *coli* and 5 μg/mL for *B*. *subtilis*), ampicillin [Amp] (100 μg/mL), or tetracycline [Tet] (2.5 μg/mL). For enumerating *C*. *difficile* titers *in vivo*, taurocholate-cefoxitin-cycloserine-fructose agar (TCCFA) containing 0.1% TA when needed, 250 μg/ml D-cycloserine, and 16 μg/ml cefoxitin^42^. All bacterial strains were stored either as glycerol stocks (16.5% v/v) at −70°C or as spores at 4°C. All bacterial strains can be found in Supplementary Table 1. Plasmids and primers used in this study can be found in Supplementary Tables 2 and 3, respectively.

### Toxin release in *C. difficile*

All strains were streaked onto BHIS supplemented with TA plates. Well-isolated colonies were picked into TY medium and grown for 16 hr. The following day, the resulting growth was washed three times in PBS to ensure minimal toxin carry-over. The strains were sub-cultured into separate 5mL inoculations with an OD_600_ of 0.01 for each time point. Aliquots were serially diluted 10-fold and spot plated onto BHIS supplemented with TA for CFU enumeration, at each time point. The cultures were centrifuged at 3200 xg for 5 min, then the supernatant was filtered through a 0.22 μM syringe filter. The pellets were washed three times in PBS. Supernatant, pellets, and whole cultures were flash frozen in liquid nitrogen and stored at −70 °C until use. Supernatants were thawed and subjected to anti-toxins ELISAs, see below. Pellets were resuspended in an equal volume of PBS with 20 μg/mL lysozyme, 1 μM pepstatin, 2 μM leupeptin, 1 mM PMSF, and 2 μg/mL DNAse. The pellets were incubated on ice for 1 hr with gentle rocking every 10 min. The pellets were sonicated (Branson Sonifier 150) using 2, 30 sec pulses. The lysates were incubated on ice for 1 min between sonication steps. The resulting cell lysates were spun down at 16,000 xg for 1 min to remove large debris. The supernatant was removed and the total protein concentration was determined using Bio-Rad Protein Assay Dye Reagent Concentrate using manufactures protocol for the standard microtiter plate protocol using BSA as the standard. Whole cultures were freeze/thawed to room temperature and snap frozen in liquid nitrogen. Two additional freeze/thaws were performed to lyse the culture.

### Anti-toxin nanobody ELISAs

Anti-TcdA nanobodies (capture: A1D8 and detection: A1C3) and anti-TcdB nanobodies (capture: B0D10 and detection B0E2) were purified as described previously^30^. ELISA plates (96-well flat-bottom; Nunc MaxiSorp) were coated overnight at 4 °C on an orbital shaker with 100 ng/mL of capture nanobody diluted in PBS. Next, the plates were washed 4 times with PBS + 0.05% Tween-20 (PBS-T), and incubated with blocking buffer (PBS-T + 2% BSA) for 2 hr at room temperature with shaking, followed by four washes with PBS-T. Supernatants, lysed pellets, and total cultures standard curves, were serially diluted (2-fold) and added to the plates. For the toxin standard curves, each toxin was diluted to 62.5 pM in blocking buffer serially diluted 2-fold. The plates were then incubated for two hours at room temperature with shaking and washed 4 times with PBS-T. Captured toxins were detected by addition of 100 μL of solutions containing biotinylated Avi-tagged detection nanobodies: A1C3 (5 ng/mL) and B0E2 (5 ng/mL). Plates were incubated for 2 hr at room temperature with shaking and washed 4 times with PBS-T. HRP conjugated Streptavidin (ThermoScientific) was diluted 1:50,000 in blocking buffer, added to the plates, and incubated for 1 hr at room temperature with shaking. Plates were washed five times with PBS-T, then 75 μl of 1 Step UltraTMB ELISA substrate solution (equilibrated to room temperature) was added to each well. ELISAs were quenched with equal volume of 2M H_2_SO_4_ and read at 450 nm in a Cytation plate reader (Biotek). The limit of detection (LOD) was defined as the lowest concentration within the linear signal range that could be distinguished from the no-toxin control as described in^43^. All recombinant TcdA and TcdB constructs were expressed in either *Bacillus megaterium* or *E*. *coli* and purified as described previously^44–46^.

### Mutagenesis of *C. difficile*

*C. difficile* mutagenesis in R20291 was performed using the CRISPR-Cas9 two-plasmid mutagenesis system, described elsewhere^47,48^. Mutagenesis in 630Δ*erm* was performed using allelic coupled exchange^41^. gDNA was extracted from *C. difficile* strains using the MasterPure Gram Positive DNA Purification Kit (Lucigen). Briefly, upstream and downstream homology arms were amplified using primers found in Supplemental Table 2. For mutagenesis in R20291, these fragments were inserted into pBL1257 (a gift from Joseph Sorg; Addgene plasmid #190481), previously digested with *Not*I and *Xho*I using Gibson assembly^49^. The resulting plasmids were digested with *Mlu*I and *Kpn*I and amplified guide gRNA were inserted into digested plasmid. For 630Δ*erm*, homology arms were inserted into pMTL-YN3 previously digested with *Asc*1 and *Sbf*1.

An ATc inducible plasmid was created by subcloning *P_tetR_* promoter from pMTL-NIC *P_tetR_* (a gift from Aimee Shen) and inserting it into pRAN737 (a gift from Craig Ellermeier) digested with *Nhe*1 and *Bam*H1 using Gibson Assembly. The resulting plasmid was digested with *Sac*1 and *Bam*H1. *tcdR* with the native ribosome binding site was amplified from R20291 gDNA. The resulting plasmid was conjugated into R20291 or 630Δ*erm* strains as described below.

All plasmids were maintained in *E. coli* DH5α and grown at 37°C, except strains harboring TcdA-mEos4a and TcdE-mScarlet which was grown at room temperature and p*P_TetR_-tcdR* which was grown at 30°C. All plasmids were whole plasmid sequenced by Plasmidsaurus using Oxford Nanopore Technology with custom analysis and annotation to ensure no off-target mutations and were transformed into *E*. *coli* pRK24.

### Plasmid conjugations into R20291 and 630Δ*erm*

pBL1256 (a gift from Joseph Sorg; Addgene plasmid #190480) was conjugated into *C*. *difficile* strain R20291 using *B*. *subtilis* strain JH BS2 described in^47^. Overnight cultures (∼ 16 hrs) of *E*. *coli* pRK24 containing the R20291 mutagenesis plasmids were grown in LB supplemented with Erm and Amp. R20291 pBL1256 grown in BHIS supplemented with Thi were back diluted 1:20 in the respective media and grown for 6 hrs. *E*. *coli* pRK24 strains were pelleted at 2,896 xg for 30 sec, supernatant removed, and the pellet was transferred to the anaerobic chamber. R20291 pBL1256 were used to resuspend the *E*. *coli* pellets (200 μl) and ∼20 μl drops were spotted onto 2 BHIS plates supplemented with Thi. After 24 hr or 48 hr of growth, the plates were scraped and resuspended in 2 mL PBS and plated onto BHIS supplemented with Thi, Kan, cefoxitin, and linomycin. Resulting transconjugates were screened for TcdB using primer pair 246 and 248 and for the plasmid using primer pairs listed in Supplemental table 2.

Overnight cultures (∼ 16 hrs, 1mL) of *E*. *coli* pRK24 containing the 630Δ*erm*Δ*pyrE* allelic exchange plasmid were grown in LB supplemented with Cam and Amp was pelleted at 2,896 xg for 30 s. The pellet was resuspended with 200 μL of overnight 630Δ*erm*Δ*pyrE* grown in BHIS and ∼20 μl drops were spotted onto one BHIS plates. After 8-10 hrs of growth, plates were scrapped and resuspended in 800 μL of PBS, 100 μL were spread onto BHIS containing Thi(10), Kan, and cefoxitin plates. After 3-4 days of growth, colonies were restruck onto fresh BHIS containing Thi(15), Kan, and cefoxitin plates. Only colonies that could form large colonies (single transconjugates) after 24 hrs, were struck onto CDDM+5-FOA+uracil to induce a double crossover. Colonies that grew after 2-4 days were screened for deletion of *tcdE*. The *pyrE* gene was restored by conjugating with pMYL-YN1C as described above, except screening for colonies that can grow on CDDM in the absence of uracil.

p*P_TetR_* plasmids were conjugated into various R20291 or 630Δ*erm* strains using similar conjugations methods described above. Overnight cultures (∼ 16 hrs) of *E*. *coli* pRK24 containing the p*P_TetR_* plasmids were grown in LB supplemented with Thi and Amp. Overnight strains of R20291 were grown in BHIS were diluted 1:20 and grown for 6 hrs. *E*. *coli* pRK24 strains were pelleted at 2,896 xg for 30 sec, supernatant removed, and the pellet was transferred to the anaerobic chamber. R20291 strains were used to resuspend the *E*. *coli* pellets (200 μl) and ∼20 μl drops were spotted onto 2 BHIS plates. After 24 hr or 48 hr of growth, the plates were scraped and resuspended in 2 mL PBS and plated onto BHIS supplemented with Thi, Kan, and cefoxitin. Overnight cultures (∼ 16 hrs, 1mL) of *E*. *coli* pRK24 containing the p*P_TetR_*plasmids was pelleted at 2,896 xg for 30 s. The pellet was resuspended with 200 μL of overnight 630Δ*erm* strains grown in BHIS and ∼20 μl drops were spotted onto one BHIS plates. After 8-10 hrs of growth, plates were scrapped and resuspended in 800 μL of PBS, 100 μL were spread onto BHIS containing Thi, Kan, and cefoxitin plates. Resulting transconjugates were screened for TcdB using primer pair 246 and 248 and for the plasmid using primer pairs listed in Supplemental table 2.

### Induction of CRISPR-Cas9 and whole genome sequencing

CRISPR was induced as previously described^47^. The strains were then validated to have the intended modifications by colony PCR and whole genome sequencing. The sequencing was performed on the Illumina NextSeq2000 platform using a 300 cycle flow cell kit to produce 2×150bp paired reads. 1–2% PhiX control was spiked into the run to support optimal base calling. Read demultiplexing, read trimming, and run analytics were performed using DRAGEN v3.10.12, an on-board analysis software on the NextSeq2000. Sequencing reads were deposited to NCBI Sequencing Read Archive accession number: 1628386.

### *C. difficile* growth curves

*C*. *difficile* strains were plated on BHIS supplemented with TA from glycerol stocks. Next day, a well isolated colony was inoculated into 2.5 mL TY and grown statically for 18 hrs. Overnight cultures were subcultured to an OD_600_ of 0.01 in TY medium and 200 μL of culture were added per well in two pre-reduced 96-well round bottom plates (Corning 3788). One plate was inserted into Stratus kinetic microplate reader (Cerillo), and readings were taken 30 min for 24 hours at 37°C. Each condition was done in technical triplicate. Readings were subtracted from the TY medium blank. The other plate was incubated at 37°C and used to determine CFU’s via serial dilution. R20291 was plated on BHIS+TA to select for vegetative and spores and 630Δ*erm* strains was plated on BHIS to select for vegetative bacteria^32^. In both strains, vegetative cells were killed following incubation in 50% ethanol for 30 min, the resulting spores were serially diluted and plated on BHIS+TA^32^. CFU’s were enumerated following a 24 hr incubation at 37°C.

### *C. difficile* spore preparation

*C*. *difficile* strains were plated on BHIS supplemented with TA plates from glycerol stocks. Next day, 2.5 mL BHIS were inoculated and grown for 16 hours at 37°C. The overnight culture (1 mL) was inoculated into 40 mL of Clospore medium which was then grown for 7 days anaerobically at 37°C ^50^. The suspension was centrifuged at 1980 xg for 20 minutes at 4°C, and the pellet was washed three times in equivolume of cold sterile water, or until the supernatant was clear. Spores were resuspended in 1 mL of sterile water and heat treated at 65°C for 20 minutes to eliminate vegetative cells. Spores were enumerated on BHIS supplemented with TA.

### Ethics statement

This study was approved by the Institutional Animal Care and Use Committee at Vanderbilt University Medical Center. Our laboratory animal facility is AAALAC-accredited and adheres to guidelines described in the Guide for the Care and Use of Laboratory Animals. Mice were housed in a pathogen-free room with 12-hour cycles of light and dark with clean bedding and free access to food and water. Cages were changed every two weeks. Prior to antibiotic treatment, mice were assimilated to the new facility for one week to reduce stress. All animal manipulations were performed in a laminar flow hood. The health of the mice was monitored daily, and severely moribund animals, including those with weight loss of more than 20% of the initial mass, were humanely euthanized. At the end of the studies, mice were humanely euthanized by CO_2_ inhalation.

### *C. difficile* mouse infection

C57BL/6J (JAX stock #664) female mice were purchased between 8–10 weeks of age from Jackson Laboratories. Mice were given 0.5 mg/ml cefoperazone (Sigma C4292) in drinking water for 5 days^44–46^. Antibiotic water was refreshed every other day to prevent antibiotic breakdown. After 5 days, mice were switched to regular water to recover for 2 days before inoculation. Mice were orally gavaged with 10^2^ *C. difficile* spores. R20291 mouse infection experiments were independently performed 2 times with a total of 15 mice per group while 630Δ*erm* were performed once with 5 mice per group. Mouse weight and symptoms were recorded daily, and stool samples were also collected daily (7 days for R20291 and 4 days for 630Δ*erm* infections).

The colon was prepared one of two ways for the R20291 infections (5 mice), the cecum was excised from each animal, and small nick was made in cecal apex so that cecal material could be gently squeezed out. For histopathology, the cecal tissue was laid flat on Whatman filter paper, and fixed in 10% neutral buffered formalin for 18 hours at 4°C.

After fixation, cecum was cut into pieces using the CecAx preservation method^51^. The colon was excised from each animal, flushed with PBS, opened longitudinally on Whatman filter paper, and fixed in 10% neutral buffered formalin for 18 hours at 4°C. After fixation, the colon was rolled into a Swiss roll. All tissues were transferred to 70% ethanol. For the 630Δ*erm* infections and remaining R20291 infections (10 mice), the cecal tissue was laid flat on Whatman filter paper, and fixed in freshly prepared Carnoy’s solution (60% ethanol, 30% chloroform, 10% glacial acetic acid) for 18 hours at 4°C. After fixation, cecum was cut into pieces using the CecAx preservation method. The colon was excised from each animal, rolled on Whatman filter paper without flushing and cutting, and fixed in 10 in freshly prepared Carnoy’s solution for 18 hours at 4°C and then transferred to 100% ethanol. All tissues were submitted to the Translational Pathology Shared Resource at Vanderbilt University Medical Center for paraffin embedding and serial sectioning.

For histopathology scoring, tissue sections were stained with hematoxylin & eosin using a staining kit (Abcam ab245880) and submitted to a board-certified gastrointestinal pathologist to blind score edema, inflammation, and epithelial damage from 0–4 as previously described^31,43^. Briefly, the following methods were used: the edema scores: 0—no edema; 1—mild edema with minimal multifocal submucosal expansion; 2— moderate edema with moderate multifocal sub-mucosal expansion; 3—severe edema with severe multifocal sub-mucosal expansion; 4—same as score 3 with diffuse sub-mucosal expansion. Inflammation scores: 0—no inflammation; 1—minimal multifocal neutrophilic inflammation; 2—moderate multifocal neutrophilic inflammation (greater submucosal involvement); 3—severe multifocal to coalescing neutrophilic inflammation (greater submucosal ± mural involvement; 4—same as score 3 with abscesses or extensive mural involvement. Epithelial damage scores: 0—no epithelial changes; 1— minimal multifocal superficial epithelial damage (vacuolation, apoptotic figures, villus tip attenuation/necrosis); 2—moderate multifocal superficial epithelial damage (vacuolation, apoptotic figures, villus tip attenuation/necrosis); 3—severe multifocal epithelial damage (same as above) +/− pseudomembrane (intraluminal neutrophils, sloughed epithelium in a fibrinous matrix); 4—same as score 3 with significant pseudomembrane or epithelial ulceration (focal complete loss of epithelium). Representative H&E images were taken using a Leica SCN400 Slide Scanner at 20x magnification (to a resolution of 0.5 μm/pixel). Whole slide imaging was performed in the Digital Histology Shared Resource at Vanderbilt University Medical Center (vumc.org/dhsr).

Daily fecal samples were resuspended in 1 mL PBS and were serially diluted and plated for vegetative and spores (R20291) or vegetative (*630*Δ*erm*) CFU’s on TFCCA or FCCA semi-selective medium, respectively, to ensure colonization. The resulting dilutions were heated to 65°C for 20 min to heat-kill vegetative bacteria. The heat-killed bacteria were plated onto TFCCA for spore CFU’s. Plates were enumerated following 24 hrs at 37°C.

### Light Microscopy

*C. difficile* strains were grown in 5 mL of TY medium for 18 hrs, with antibiotics or ATc (20 ng/mL), if needed. ATc induction occurred for 4 hrs. Samples were centrifuged for 5 min at 3000 xg at 4°C, and washed with an equal volume of PBS. The resulting pellet was suspended in CellBrite®Fix (Biotium) diluted 1:1000 in 1 mL of PBS for 20 min at 25°C in the dark. Following membrane labeling, the cells were washed three times in PBS and fixed in 1% para-formaldehyde (PFA) for 10 min, aerobically. The cells were washed an additional three times in PBS to remove residual PFA. The bacteria were immediately mounted onto plasma cleaned cover slips (FisherBrand #1, 22 mm) containing Prolong Gold Anti-fade mounting medium (without DAPI) (Molecular Probes). The slides were store at 4°C in the dark until imaged.

Images were recorded using a Leica Stellaris White Light Laser Confocal Microscope 63x/1.40 Oil objective at a stage temperature of 25 °C. The Leica Application Suite X (LAS X) software platform was used for acquisition and the resulting images were analyzed using Fiji and GraphPad Prism^52^. Laser intensities and detector gain were matched between samples for each experimental condition.

### Freeze-substituted electron microscopy

*C. difficile* R20291 strains containing p*PtetR* plasmids were grown for 18 hrs and induced with ATc for 4 hrs and was three times in PBS. Cells were fixed in 2.5% glutaraldehyde for 30 min at room temperature followed by overnight incubation at 4°C. Fixed pellets were loaded into 3 mm high pressure freeze hats and vitrified using a Leica EM ICE high-pressure freezer. The samples were directly transferred to a Leica AFS2 freeze substitution device using 1% uranyl acetate in methanol at −90°C for 48 hr and then gradually warmed to −45°C over the course of 12 hr. The samples were infiltrated with 50%, 75%, and 100% HM20 lowicryl for 4 hr each using methanol as the transition solvent. The resin was UV polymerized at −45°C for 24 hr followed by 0°C for 48 hr. Samples were sectioned at a nominal thickness of 200 nm on a Leica Enuity Ultramicrotome. Samples were imaged on a JEOL 2100 at 200 keV using an AMT nanosprint15mkII CMOS camera. SerialEM was used for data collection.

### Plunge freezing for cryo-ET imaging

R20291-*tcdA-mEos4a* containing p*P_tetR_-tcdR* was grown for 18 hrs and induced with ATc for 4 hrs. The cells were fixed in 1% PFA as described previously. Following fixation, the bacteria were diluted to an OD_600_ ranging from 1-10. BSA 10 nm gold tracers (60 µL of OD_552_ of 1) were washed three times with PBS. Equal volume of bacteria were used to resuspend the gold fiducials. Quantifoil LondonFinder R3.5×1 200 mesh grids were glow discharged using a GlowCube for 30 sec. The grid was blotted twice using a Leica GP2 at room temperature with 70% humidity for 3 sec. For each blot, 5 µL of the bacterial/fidicial mixture was added to the front of the grid and 3.5 µL of PBS was applied to the back. The resulting grids were plunged into liquid ethane and stored in liquid nitrogen until loaded into the microscope. Plunge frozen grids were clipped into FIB-autoloader grids (ThermoFisher) and were screened using the ThermoGlacios cryo-TEM at 200kV with a Falcon4 direct electron detector for ice conditions and grid orientations before cryo-confocal imaging.

### Cryo-confocal

Vitrified grids were loaded into a Stellaris cryo-shuttle and imaged on a Leica Stellaris outfitted with a EM Cryo CLEM cryogenic stage and cryo-50x objective. Grids were imaged using Leica CORAL CLEM software. Three-dimensional tilesets of mEos4a and Cellbrite 640 were acquired over the millable area of each grid, the center of grid along with holes or finder numbers were located in each tileset. Samples were stitched and exported for correlation with cryo-FIB and cryo-TEM images.

### Cryo-FIB milling

The stellaris shuttle was directly transferred to a Leica pretilted CLEM holder for a Leica VCT500 cryo-transfer system and were coated with a 3 nm platinum using a Leica ACE600 cryo-ebeam system. Grids were transfer to a Zeiss Crossbeam 550 FIB-SEM equipped with a Leica VCT copper-band cooled cryo-stage, the gas injection system was used to deposit an organometallic platinum onto each grid for 10 sec at a working distance of 5.5 mm.

Identification of suitable targets was done by correlating the tilesets acquired in the Stellaris with the SEM (2 kV, X 20 pA) and FIB images (30 kV, X 20 pA) with the samples normal to the SEM column. Images were correlated using the BigWarp plugin for FIJI^53^. Suitable areas for milling were marked with trenches at (700 pA, 30 kV) at an incident angle of 36°. After the trenches were made the samples were tilted to a final incident angle of 8°. Lamella width was set to 15 µm with a target thickness of less than 200 nm. In total, five FIB currents were used, gradually reducing with lamella thickness from 700 pA, 300 pA and 100 pA, 50 pA, and 20 pA for final polishing. All milling was performed using Zeiss SmartFIB software using drift correction. After milling overview images of the lamella were taken normal to the SEM for further correlation in cryoTEM.

### Cryo-ET collection, reconstruction, and data processing

CryoET data were collected on ThermoTitan Krios G4 microscopes operating at 300 kV with a 100 μm objective aperture and equipped with a Gatan’s BioQuantum energy filter (slit width 10 eV), combined with a Gatan K3 direct electron detector. Data were acquired using were acquired using ThermoFisher Tomo5 software. Tilesets acquired on the Stellaris were correlated to the search maps using BigWarp plugin for FIJI^53^. Tilt series on lamellae were either recorded in a dose-symmetric tilt scheme ±45° with 2° angular increments. All tilt series on lamellae were recorded with a pixel size of 2.6 Å, a defocus of −6 μm, and had a cumulative electron dose of 125 e^−^ Å^−2^. The resulting images were analyzed in the IMOD package and 4x-binned cryo-tomograms were reconstructed by weighted back projection in IMOD^54–57^. The resulting cryo-tomograms were denoised using CryoSamba^58^. Segmentation of the inner membrane, peptidoglycan, s-layer, vesicles, and electron dense feature were performed in Segmentation Wizard of Dragonfly using the Pre-trained Deep Learning U-Net model.

### Freeze Fracture

*C. difficile* p*P_tetR_*was grown for 18 hrs and induced with ATc for 4 hrs. The cells were fixed in 1% PFA as described previously. Samples were vitrified using a Leica EM ICE high pressure freezer using domed freeze fracture planchettes. Samples were fractured at −110°C in a Leica ACE 600 freeze-fracture device, briefly etched at −100°C for 2 minutes then shadowed with 3 nm platinum at 30° and stabilized with carbon at 90° by ebeam deposition. The samples were then loaded in the Zeiss Crossbeam 550 FIBSEM and imaged by SEM (2 kV, X 50 pA) using Zeiss SmartSEM software.

### Vesicle preparation

*C. difficile* strains were grown for 18 hr in TY medium (200 mL) at 37°C. The cells were spun at 3,200 xg for 20 min at 4°C to remove vegetative bacteria and spores. The supernatant was removed and filtered using a Stericup, Vacuum Driven Disposable Filtration System with a 0.22 µm PVDF filter. The resulting filtrate was concentrated using Amicon Ultra 100K molecular weight cutoff centrifugal filters at 2,000 xg in 10 min increments to ∼50 mL, and transferred to 38×102 mm polycarbonate thick well bottles and centrifuged in a Beckman Ti45 rotor at 100,000 xg for 2 hr at 4°C. The supernatant was removed and the pellet was washed with 1 mL 25 mM MES pH 6.2 containing 1 µM Vybrant DiO (Invitrogen). The insoluble fraction was added to the top of an Optiprep Iodixonol gradient ranging from 10% to 35% in 25 mM MES pH 6.2 in a 14×89 mm ultra-clear open cap tubes. The gradients were centrifuged at 120,000 xg in a Beckman SW60 rotor for 16 hr at 4°C. Individual 1 mL fractions were isolated and tested for TcdA and TcdB using the ELISA described above, except samples were diluted 1:2 in PBST+BSA with vigorous pipetting to lyse the resulting extracellular vesicles. Fractions containing TcdA and TcdB were filtered sterilized using a Millex-GV 0.22 µm PVDF membrane filter diluted to 10 pM and added to Vero-GFP cells as described above. Fraction 6 or 9 were negatively stained for presence of EVs.

### Negative stain electron microscopy of vesicles

Carbon (300 mesh) (Electron Microscopy Sciences) grids were coated with 0.01% poly-D-lysine (Sigma) for 15 min at room temperature and the excess of the liquid was wicked off. Grids were glow discharged on low for 30 sec using a PDC-32G (Harrick Plasma). The grids were floated on 5 µL of sample for 5 min, followed by one quick wash (< 1sec) and a 10 sec wash in water. The grids were negatively stain with 2% (w/v) uranyl acetate. The negative stain was blotted by pressing the side of the TEM grid against a freshly torn piece of #1 Whatman filter paper until the grid was dry. TEM imaging was performed on a JEOL 2100 Plus operating at 200 keV using an AMT nanosprint CMOS camera. Random fields of EVs were imaged over the entire grids.

### Cell rounding assays

Vero-GFP were made as described previously^47^. The cells were cultured in Dulbecco’s modified Eagle’s medium (DMEM), with high glucose, sodium pyruvate, L-glutamine (Gibco), 10% fetal bovine serum (FBS, Corning), and 10 µg/ml puromycin at 37°C with 5% CO_2_. Vero-GFP cells were seeded into a 96-well plate at 25,000 cells per well or for a 384 well plate 5,000 cells per well in complete DMEM. After growing cells for 24 hrs, media from wells were removed and replaced with 90 µL with complete DMEM containing 10 µL of 1 pM toxin associated vesicles determined by ELISA, 10 µL DMEM or 10 µL of 100 nM anti-toxin cocktail. rTcdA and rTcdB (1 pM) were used as positive control. After intoxication, brightfield and fluorescence images were collected every 45 minutes on a Cytation 5 imager (BioTek) with an atmosphere of 5% CO2 at 37°C.

### Cell Rounding Quantification

Images acquired from cell rounding experiments were analyzed using an automated cell rounding analysis pipeline developed in CellProfiler (v4.2.1) using machine learning cell segmentation models from Cellpose (v2.0.5) and cell rounding classifiers generated in CellProfiler Analyst (v3.0.4)^59–61^. From these images, the total number of rounded and non-rounded cells were counted as described in ^62,63^. These CellProfiler pipelines and the CellProfiler Analyst model are available on https://github.com/kochild/Cell-Rounding-Pipelines-and-Models.

### Statistical analysis

All statistical tests were performed using GraphPad Prism 10 software. For statistical comparison of two groups with one independent variable, unpaired t-test, Mann-Whitney test, or lognormal Welch’s t-test were used to calculate statistical significance. For statistical comparison of two groups with two independent variables, two-way ANOVA with a Dunnett’s post hoc analysis, a two-way ANOVA followed by an Uncorrected Fisher’s LSD post hoc test, or mixed-effects analysis with Šídák’s multiple comparisons tests were used to calculate significance. The Log-rank (Mantel-Cox) multiple comparison test was used for survival curve comparisons. A *P*-value of <0.05 was considered statistically significant, with \**P*< 0.05, \*\**P*< 0.01, \*\*\**P*< 0.001, \*\*\*\**P*< 0.0001.

## Acknowledgments

The Translational Pathology Shared Resource is supported by NCI/NIH Cancer Center Support Grant P30CA068485 and the Vanderbilt Digestive Disease Research Center is supported by P30DK058404. This work was supported by the following NIH equipment awards: S10OD030292, R24OD037694. The authors would like to thank: Mariam Haider for her assistance with the Krios, Christopher Peritore-Galve for his assistance with whole genome sequencing, Tanner Durst for his assistance with counting TcdE positive bacteria, and Rachel Hart for her assistance with FIB-milling and electron microscopy sample preparations. We also thank Joe Sorg, Aimee Shen, and Craig Ellermeier for sharing plasmids and strains. This work was supported by NIH Ruth Kirschstein Award F32GM139303 (SLK), VA CDA BX005699 (NOC), AI174999 (DBL), BX002943 (DBL), and AI095755 (DBL).

## Extended Data Figures

**Extended Data Figure 1. Assessing the role of *tcdE* on toxin release in *C. difficile*.** Bacterial cultures were grown in TY medium and at various time points TcdA and TcdB were measured by ELISA. **A,D)** Culture TcdA or TcdB measured in R20291 (black) and R20291Δ*tcdE* (pink) at 4 hrs (mid log phase), 8 hrs (early stationary phase), and 18 hrs (death phase) time post inoculation. **B,C,E,F)** Supernatant and pellet data are the same as shown in Figure 2. **G,J)** Culture TcdA or TcdB measured in 630Δ*erm* (black), 630Δ*erm*Δ*tcdE* (pink), at 4 hrs (log phase), 8 hrs (early stationary phase) and 18 hrs (late stationary phase) time post inoculation. **H,I,K,L,F)** Supernatant and pellet data are the same as shown in Figure 2. An unpaired t-test was used to calculate statistical significance (* *P*<0.05, ** *P*<0.01). Empty circles represent ELISA values that fell below the limit of detection, dotted line. Each data point is the average of two technical replicates. Error bars represent SEM.

**Extended Data Figure 2. The deletion of TcdE in R20291 results in attenuated weight loss and reduced disease severity in a mouse model of CDI.** Mice were infected by oral gavage with 100 spores of **(A)** R20291, **(B)** R20291Δ*tcdE*, or **(C)** PBS vehicle control. Weight loss was tracked for each individual mouse for 7 days. Dotted line indicates humane endpoint. Arrows indicate time point where mice were euthanized. **(D)** Percent weight change from day 0 of infection study. The weight of each mouse was collected daily and plotted as relative weight change from day 0. A mixed effects analysis with a Dunnett’s post hoc analysis was used to determine significance (** *P*<0.01, *** *P*<0.001, **** *P*<0.0001). Cecal **(E,G)** and colonic **(F,H)** tissues were harvested at day 4 post infection, stained with H&E, and scored for edema (stars), inflammation (triangles) and epithelial injury (arrows). Scale bars represent 100 µm. A Mann Whitney U test was used to determine significance (* *P*<0.05, ** *P*<0.01). Average shown and error bars represent SEM.

**Extended Data Figure 3. Stool from mice infected with R20291**Δ***tcdE* contains similar amounts of TcdA and TcdB compared to R20291.** Mice were infected with 100 spores of R20291 (black) or R20291Δ*tcdE* (pink) and disease was tracked for 7 days. Stool was collected every day for 7 days, and anti-toxin ELISAs were performed. Stool was diluted to 10 mg/mL for **(A)** TcdA or 0.5 mg/mL for **(B)** TcdB. Average is shown and error bars represent SEM. Dotted line represents limit of detection, samples that are empty circles represent values that fall below limit of detection. Šídák’s multiple comparison tests was used to calculate statistical significance (*** *P*<0.001).

**Extended Data Figure 4. The deletion of TcdE in 630**Δ*erm* does not affect weight loss or disease severity in a mouse model of CDI. Mice were infected with 100 spores of **(A)** 630Δ*erm* (black) or **(B)** 630Δ*erm*Δ*tcdE* (pink). Weight loss was tracked for each individual mouse for 4 days. Dotted line indicates humane endpoint. Arrows indicate time point where mice were euthanized. **C)** Percent weight change from day 0 of infection study. The weight of each mouse was collected daily and plotted as relative weight change from day 0. Average is shown and error bars represent SEM. A mixed effects analysis with a Dunnett’s post hoc analysis was used to determine significance. Cecal **(D,F)** and colonic **(E,H)** tissues were harvested at day 4 post infection, stained with H&E, and scored for edema (stars), inflammation (triangles) and epithelial injury (arrows). Scale bars represent 100 µm. A Mann-Whitney U test was used to determine significance. Average shown and error bars represent SEM.

**Extended Data Figure 5. Stool from mice infected with 630**Δ***erm***Δ***tcdE* contains similar amounts of TcdA and TcdB compared to 630**Δ***erm*.** Mice were infected with 100 spores of 630Δ*erm* (black) or 630Δ*erm*Δ*tcdE* (pink) and disease was tracked for 7 days. Stool was collected every day for 7 days and anti-toxin ELISAs were performed. Stool was diluted to 10 mg/mL for **(A)** TcdA or 0.5 mg/mL for **(B)** TcdB. Average is shown and error bars represent SEM. Dotted line represents limit of detection, samples that are empty circles represent values that fall below limit of detection. Šídák’s multiple comparison tests was used to calculate statistical significance.

**Extended Data Figure 6. TcdE contributes to cellular lysis in R20291.** The growth dynamics of R20291 or R20291Δ*tcdE* containing empty vector (p*P_tetR_,* yellow) or anhydrous tetracycline inducible *tcdR* (p*P_tetR_-tcdR*, orange) were followed by optical density and plating. **A,C)** Optical density was read every 30 min for 24 hrs. **B,D)** The mixture of vegetative bacteria and spores or spores alone were plated every 2, 4, 6, 8 and 24 hrs. R20291 or R20291Δ*tcdE* containing p*P_tetR_* (yellow) or p*P_tetR_-tcdR* (orange) induced with 20 ng/mL anhydrous tetracycline (ATc) **E,G)** Optical density was read every 30 min for 24 hrs. **F,H)** Vegetative and spores or spores alone were plated every 2, 4, 6, 8 and 24 hrs. Multiple lognormal Welch’s t test was used to determine significance (\**P*<0.05, \*\**P*<0.01). Data is shown as average, and error bars represent SEM. Dotted line represent limit of detection. Empty circles (p*P_tetR_*) or squares (p*P_tetR_-tcdR*) represent values that fell below limit of detection.

**Extended Data Figure 7. TcdE contributes to cellular lysis in 630**Δ***erm*.** 630Δ*erm* or 630Δ*erm*Δ*tcdE* containing empty vector (p*P_tetR_,* yellow) or anhydrous tetracycline inducible *tcdR* (p*P_tetR_-tcdR*, orange). **A,C)** Optical density was read every 30 min for 24 hrs. **B,D)** Vegetative bacteria or spores were plated every 2, 4, 6, 8 and 24 hrs. 630Δ*erm* or 630Δ*erm*Δ*tcdE* containing p*P_tetR_*(yellow) or p*P_tetR_-tcdR* (orange) induced with 20 ng/mL anhydrous tetracycline (ATc) **E,G)** Optical density was read every 30 min for 24 hrs. **F,H)** Vegetative or spores were plated every 2, 4, 6, 8 and 24 hrs. Multiple lognormal Welch’s t test was used to determine significance (\**P*<0.05, \*\**P*<0.01). Data is shown as average, and error bars represent SEM. Dotted line represent limit of detection. Empty circles (p*P_tetR_*) or squares (p*P_tetR_-tcdR*) represents values that fell below limit of detection.

**Extended Data Figure 8. Overexpression of tcdR causes increased membrane production resulting in vesicle formation and cell lysis**. Freeze-substituted transmission microscopy micrographs of R20291 or R20291Δ*tcdE* empty vector control **(A)** alone or **(B)** with induction of *tcdR* by *P_tetR_* (scale bar 500nm). **C)** Tomogram of R20291 overexpressing *tcdR* (scale bar 500nm).

**Extended Data Figure 13. R20291 EVs contain TcdA and TcdB and are capable of rounding Vero-GFP cells.** EVs from strains **(A)** R20291, **(B)** R20291Δ*tcdE,* and **(C)** R20291 *tcdAB::CT* were isolated, concentrated, and crude membrane fractions were purified using ultracentrifugation. The resulting pellet was applied to an iodixanol gradient. Anti-TcdA and TcdB ELISAs were used to measure toxin concentrations in each fraction. Boxes represent fractions used for negative stain electron microscopy. **D)** Vero-GFP cells were used in cell rounding assays. 10 pM rTcdA/TcdB, 10 pM of EVs from fractions 9-11, and 10 µL in the absence of toxin were added to the cells. Anti-TcdA/B anti-toxin was added when noted. Images were captured every 30 minutes for 3 hrs. **E)** The resulting images were segmented to determine percent cell rounding. (Scale bar 500nm).

**Supplemental Movie 1.** Fully segmented 3D tomogram from Figure 5A. Segmented tomogram overviews represent inner membrane, red; peptidoglycan, yellow.

**Supplemental Movie 2.** Fully segmented 3D tomogram from Figure 5F. Segmented tomogram overviews represent inner membrane, red; peptidoglycan, yellow; S-layer, blue; electron dense region, green; lipid bilayer vesicles, purple.

