## Extended data figures for "A targeted cell lysis mechanism facilitates toxin release in Clostridioides difficile"

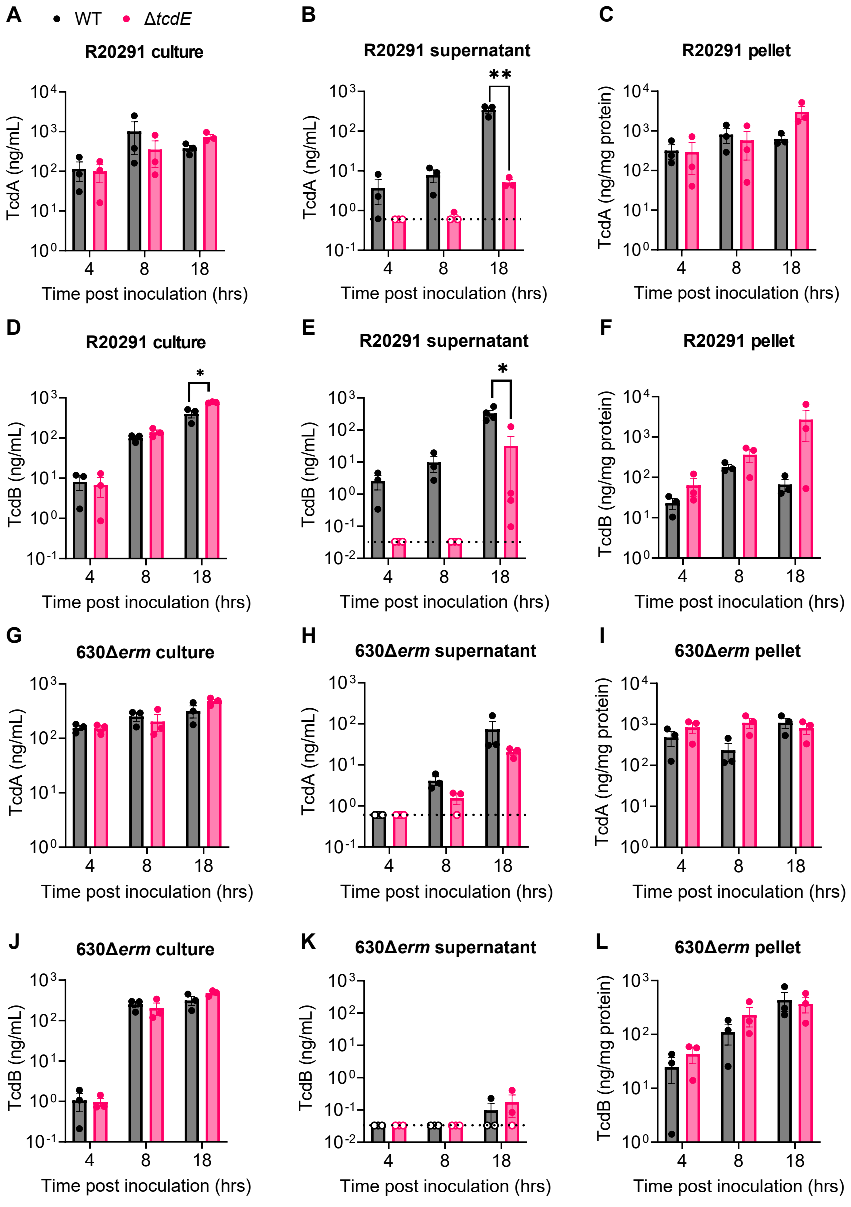


**Extended Data Figure 1.**

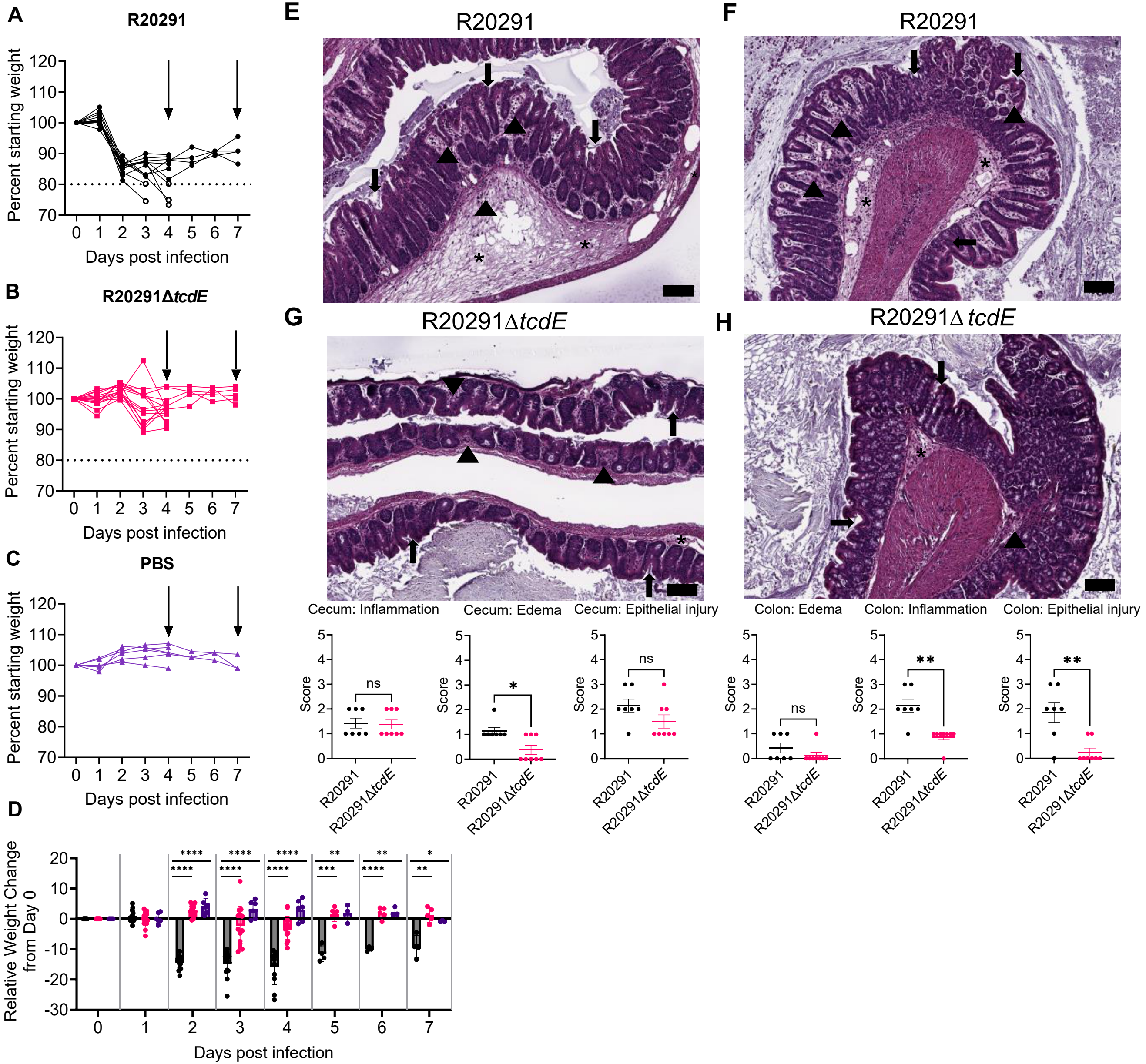


**Extended Data Figure 2.**

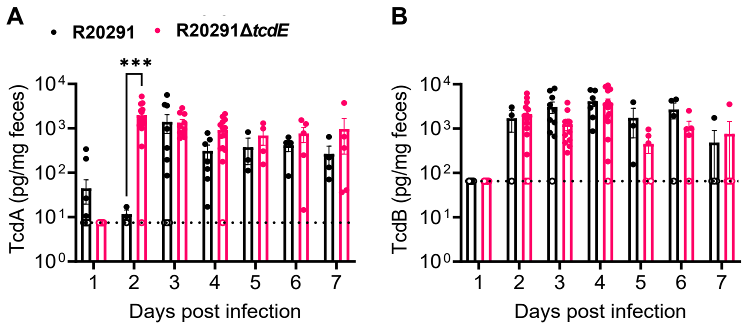


**Extended Data Figure 3.**

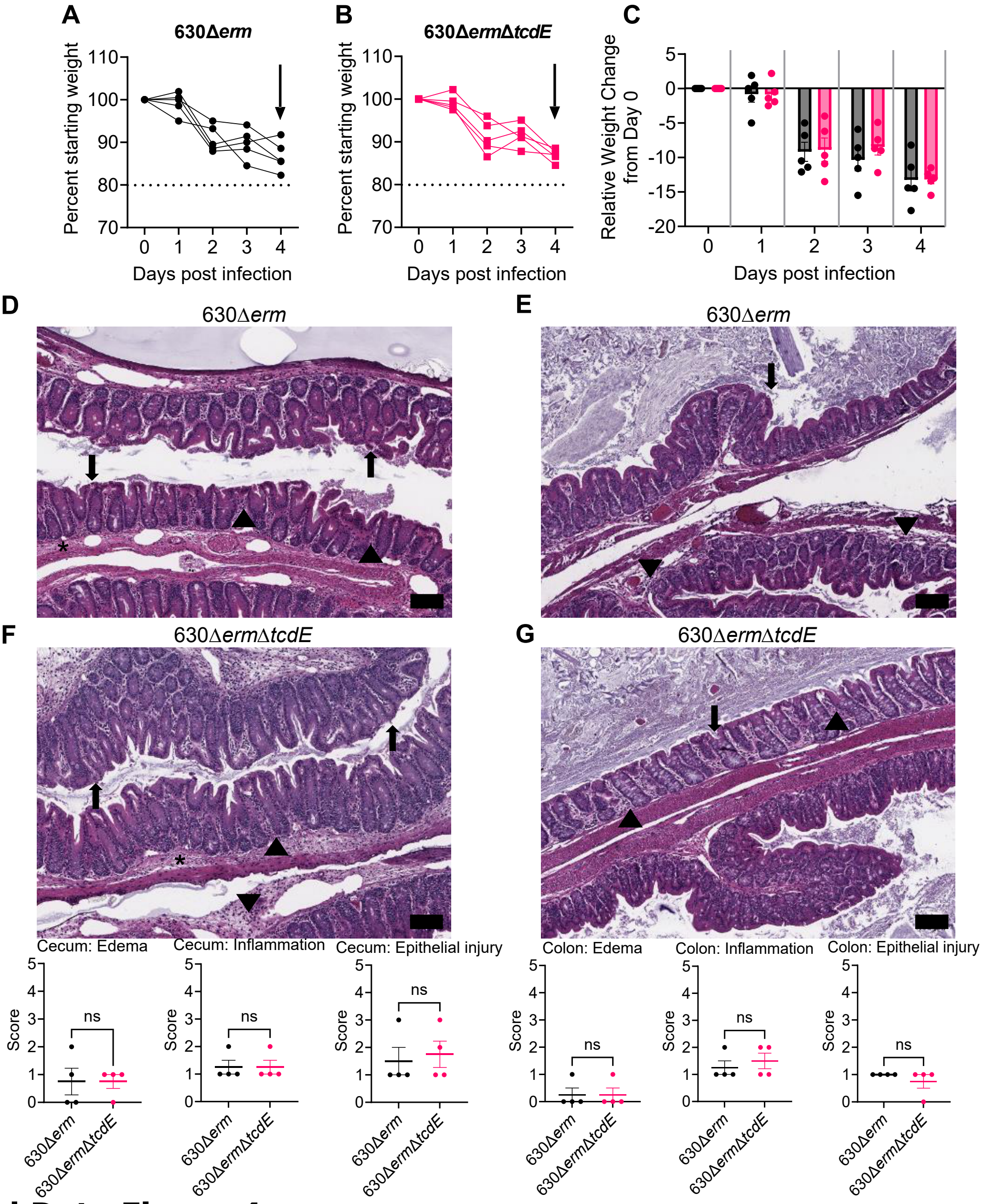


**Extended Data Figure 4.**

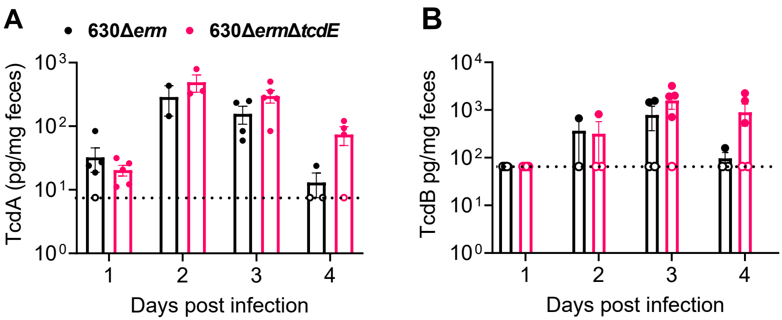


**Extended Data Figure 5.**

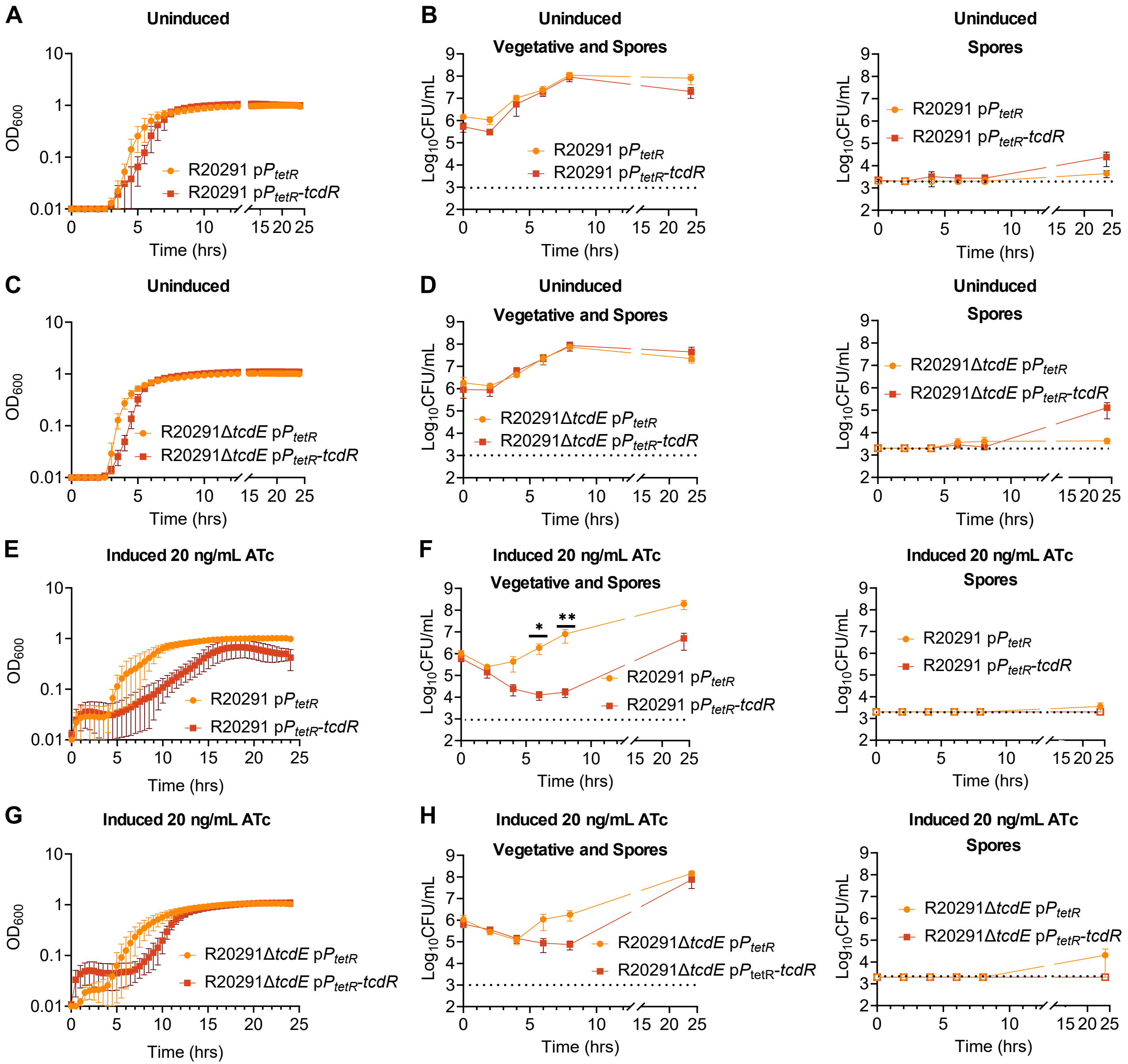


**Extended Data Figure 6.**

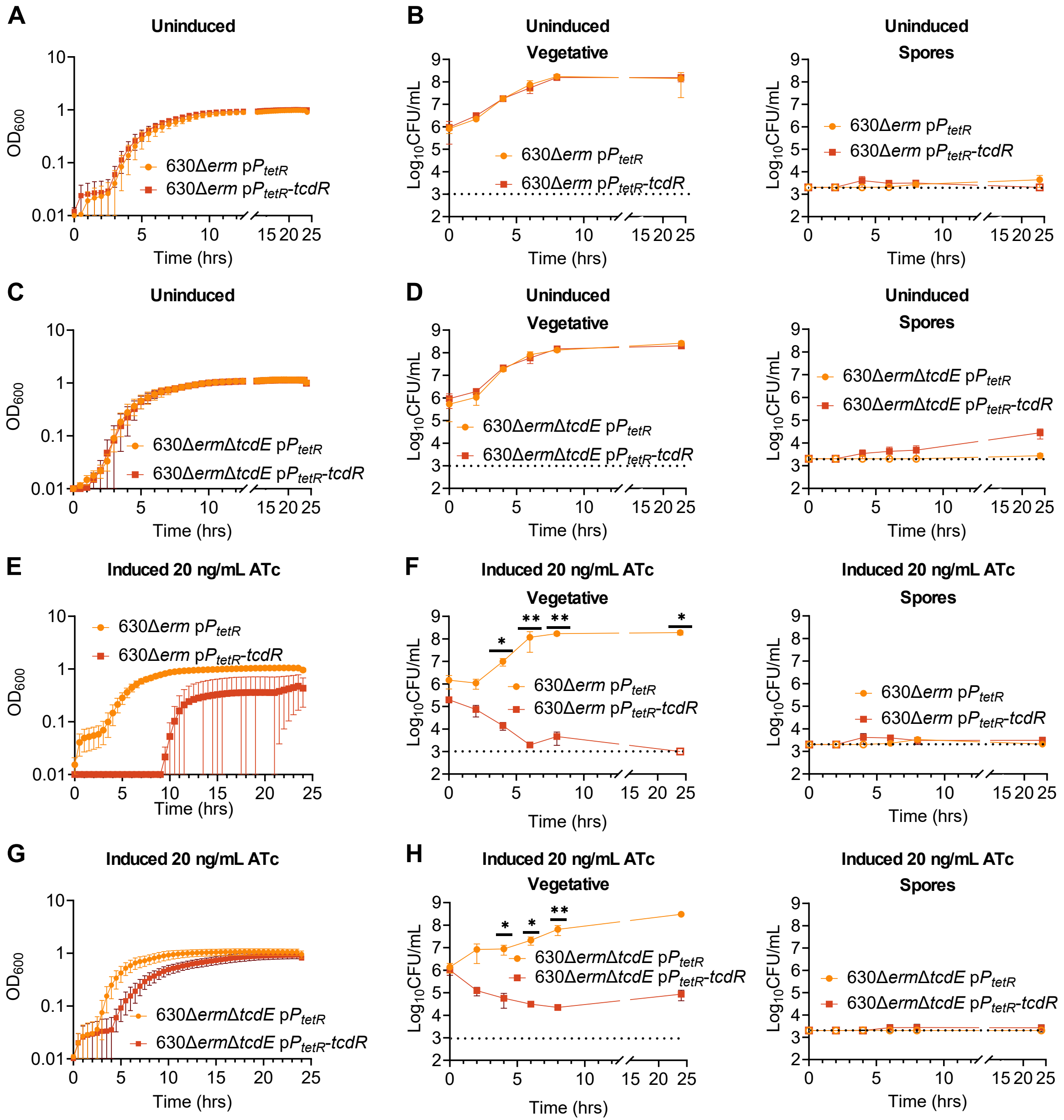


**Extended Data Figure 7.**

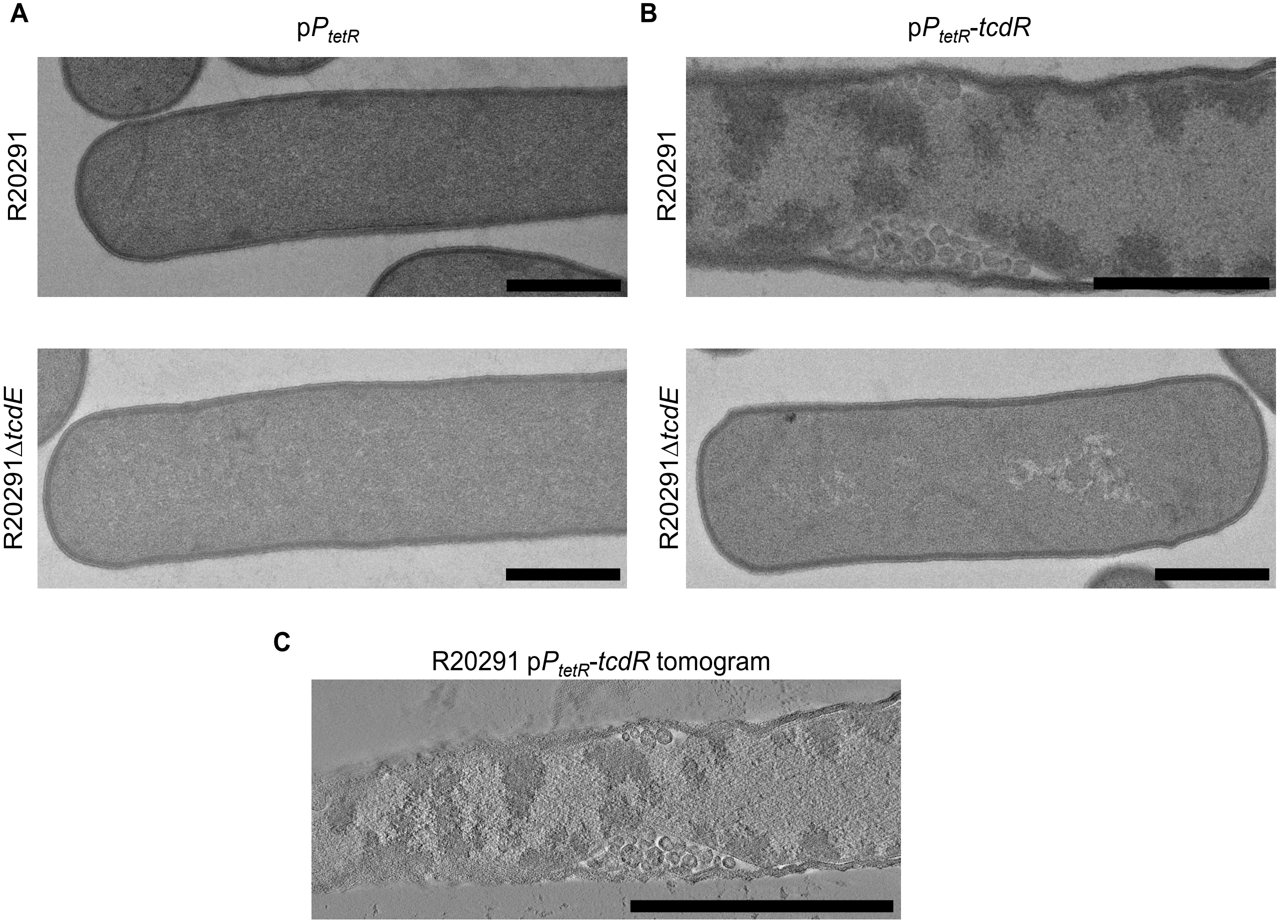


**Extended Data Figure 8.**

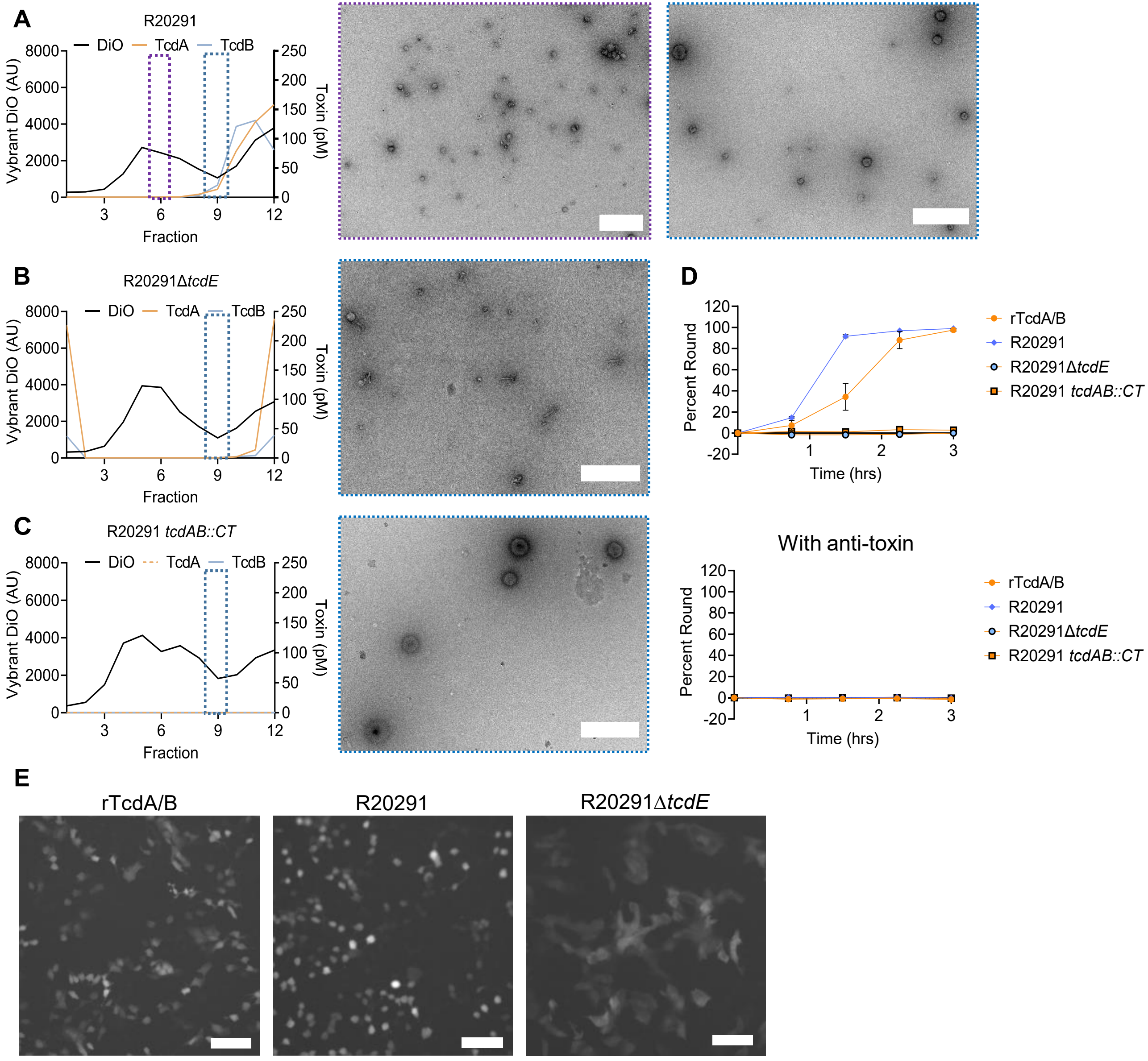


**Extended Data Figure 9.**
